# Forfeiting the founder effect: turnover defines biofilm community succession

**DOI:** 10.1101/282574

**Authors:** Colin J. Brislawn, Emily B. Graham, Karl Dana, Peter Ihardt, Sarah J. Fansler, William B. Chrisler, John B. Cliff, James C. Stegen, James J. Moran, Hans C. Bernstein

## Abstract

Microbial community succession is a fundamental process that effects underlying functions of almost all ecosystems; yet the roles and fates of the most abundant colonizers are poorly understood. Does early abundance spur long term persistence? How do deterministic and stochastic processes influence the roles of founder species? We performed a succession experiment within a hypersaline microbial mat ecosystem to investigate how ecological processes contributed to the turnover of founder species. Bacterial and micro-eukaryotic founder species were identified from primary succession and tracked through a defined maturation period using 16S and 18S rRNA gene amplicon sequencing in combination with high resolution imaging that utilized stable isotope tracers to evaluate basic functional capabilities. The majority of the founder species did not maintain high relative abundances in later stages of succession. Turnover (versus nestedness) was the dominant process shaping the final community structure. We also asked if different ecological processes acted on bacteria versus eukaryotes during successional stages and found that deterministic and stochastic forces corresponded more with eukaryote and bacterial colonization, respectively. Our results show that taxa from different kingdoms, that share habitat in the tight spatial confines of a biofilm, were influenced by different ecological forces and time scales of succession.

## INTRODUCTION

While each microbial community is distinct and subject to specific environmental limitations and opportunities, some generalized principles guide community assembly and succession (Nemergut et al 2013). For example, the arrival or disappearance of species exert feed-back responses onto the local habitat. The niches available for microbial life strategies can change during community succession for every new species and/or function recruited (Jones et al 1997). Another generalized principle is that almost all microbial communities studied to date tend toward species abundance distributions where the majority of the taxa are found in low relative abundance (Curtis et al 2002, Delgado-Baquerizo et al 2018, Magurran 2013). Some evidence suggests that the minority of taxa found in high abundance correlate better with environmental factors than complete microbiomes (Gobet et al 2010). These findings lend credibility to the idea that abundant members apply a greater influence on their environment and thus, influence the recruitment and selection of other taxa – *i.e.*, exert priority effects (Fukami et al 2010). The ecological processes underlying the fate of colonizers found in high abundance (founder species) is often unexplored. Does early abundance relate to persistence or are founder species turned over as a microbiome matures and establishes new selection forces?

In this context, turnover is an ecological process that accounts for the processes by which existing species are replaced with incoming ones and implies that new taxa are introduced into a given spatial domain via dispersal (Baselga 2012). Another ecological process that drives the distinction of a community’s structure from its different observable successional states can be described by nestedness, which accounts for species loss without replacement. Here, we apply the nestedness concept as it has been used before to describe those parts of a community that are nested subsets of its prior taxonomic composition. We used the combination of nestedness and turnover to account for the beta diversity observed between communities at different periods in time (Baselga et al 2007, Legendre and Gauthier 2014, Whittaker 1960). We were curious if different selective pressures or patterns of succession could be observed between bacterial and micro-eukaryotic members of the microbiome. This biofilm succession experiment was therefore study to provide new knowledge about how microbes belonging to different kingdoms are influenced by different ecological forces and timescales of community succession.

Due to the near ubiquity and extreme functional diversity of microbial communities, generalized principles that relate a microbiome’s history to its current state remain elusive. Much remains unknown about the roles that colonizing microbes play in influencing the future states of their respective microbiomes. Hence, microbial community succession remains an active area of research (Castle et al 2017, Knelman et al 2018, Kolinko et al 2018). We do know, however, that these processes can be influenced by both deterministic and stochastic factors (Nemergut et al 2013, Vellend 2010, Zhou and Ning 2017). Deterministic ecological processes include selection or environmental filtering, which are governed by the limitations and opportunities imposed by both the environment and niches created by predecessors. Presumably, the founder species play the largest role in establishing the trajectory for succession because of the assumed relationship between high abundance and environmental influence. However, stochastic processes may also be strong influencers. This is especially true when selection is weak (Chase and Myers 2011, Stegen et al 2012). We posit that if the combined effects of founder species are able to strongly influence their localized environment (i.e., impose strong selection) then they will more strongly direct recruitment of new taxa during community succession (Fukami et al 2010). This effect is expected to be manifested in a high degree of phylogenetic relatedness in the successful incoming species. This postulate comes with an assumption that related taxa are more likely to be recruited together because they are expected to have similar habitat associations. While the validity of this assumption can be debated, multiple studies have shown that relationships between phylogenetic relatedness, habitat selection and functional traits are observable (Andersson et al 2010, Bryant et al 2008, Martiny et al 2013, Wang et al 2013).

We performed a microbial community succession study in a standardized 79-day colonization experiment and placed specific emphasis on the founder species as the community matured. This was done *in situ* within the unique Hot Lake ecosystem, which is a MgSO_4_-dominated hypersaline microbial mat located in North Central Washington State (USA). The microbial ecosystem was chosen because of *a priori* knowledge of seasonal community development presented in a previous study that ascertained a distinct mat-development period during summer months (June-August) (Lindemann et al 2013). The microbiome characterized from the mat during this growth period remained remarkably similar, in terms of members and relative abundance of taxa, across multiple related studies (Bernstein et al 2017a, Bernstein et al 2017b, Mobberley et al 2017, Moran et al 2014). Our first goal was to determine the structure and function (autotrophy and diazotrophy) of the community from the colonization period through to the season’s typical maturation period. Next, we quantified the relative contributions of turnover and nestedness to the observable beta diversity across this same time scale. We used a high degree of sample replication (n = 20) in combination with ecological null models commonly used to infer the relative influences of stochastic and deterministic forces at each period of succession. Lastly, we identified the founder species and determined how their abundances changed from the colonization through the maturation periods. We found relatively few highly abundant colonizers that were defined as founder species and of these, most were outcompeted in later stages of succession. We then determined that turnover (versus nestedness) was the dominant process that shaped the final structure of the mature community. Along with turnover and nestedness, the relative contributions of deterministic and stochastic ecological forces varied with respect to the bacterial and eukaryotic components of the microbiome. Null model analyses indicated that the bacterial components of the community trended towards homogenous selection and that the bacterial founder species likely modified their localized environment in ways that mediated environmental filtering of other bacteria. In contrast, colonization of eukaryotic founder species corresponded with weak variable selection and trended towards stochastic processes. These results highlight the fact that although species from different kingdoms are often tightly entangled within the same microbial habitat, the respective forces that shape their community dynamics can be distinct.

## MATERIALS AND METHODS

### *In situ* incubation and Sampling

We deployed sterile, glass substrata in Hot Lake, which is a MgSO_4_ dominated, hypersaline lake in north-central Washington State (USA); 48.973062°N, 119.476876°W (Fig. S1) (Anderson 1958, Moran et al 2014, Zachara et al 2016). The surfaces included plain borosilicate microscope slides (75 × 25 mm) and round class coverslips (1 in. diameter) placed in large glass petri dishes (15 cm diameter). Each dish contained either 5 slides or 15 coverslips; 55 dishes were deployed totaling 205 slides and 210 coverslips. Incubations were initiated at 13:00 May 20^th^, 2015 when all surfaces were place on the bottom of the lake at an initial depth of 43-50 cm. One day prior to deployment, the native mat was denuded from the incubation site. The first sample was taken at 16:00 May 20^th^, 2015 (2 h of incubation); the following samples were extracted on near 12:00: May 28^th^, June 3^rd^, June 24^th^, July 15^th^ and August 6^th^ (2015). Samples (at least 30 slides and coverslips per time point) were removed from the lake using an inflatable raft to avoid disturbing the sediment; lake water was simultaneously collected from the sampling position; each slide was washed in the native water to remove loose sediment using sterile forceps. Samples were fixed in the field for the following: gDNA extraction (20 slides per time point; stored on dry ice); confocal imaging (5 slides per time point; fixed in 4% paraformaldehyde; stored at 4°C); stable isotope probing for nano-SIMS imaging (15 coverslips per time point; fixed in 4% paraformaldehyde stored at 4°C).

### Amplicon sequencing

gDNA was extracted using the MoBio PowerSoil DNA isolation kit (Qiagen, Carlsbad, CA) in accordance with the Earth Microbiome Project (EMP) protocols (Gilbert et al 2010). Sequencing was performed on an Illumina MiSeq instrument (Illumina, San Diego, CA). Twenty separate 16S and 18S rRNA gene amplification reactions were performed on template from each extraction. The extraction and amplification were successful for 4 slides from day-1, 5 slides from day-8 and all 20 slides for every other sample. The 16S primers targeted the V4 hypervariable region of the 16S SSU rRNA gene using the V4 forward primer (515F) and V4 reverse primer (806R) with 0–3 random bases and the Illumina sequencing primer binding site (Caporaso et al 2010). The 18S primers targeted the V9 hypervariable region of the 18S SSU rRNA gene (Amaral-Zettler et al 2009).

### Amplicon analysis

Illumina reads were processed with VSEARCH 2.0.2, an open-source tool for search and clustering (Caporaso et al 2010, Rognes et al 2016); overlapping 16S reads were paired, filtered to a maximum expected error of 1 bp per read, and labeled. Reads were pooled, de-replicated, and chimera-checked with the VSEARCH followed by UCHIME-ref using the RDP Gold database (Wang et al 2007). After discarding chimeras and singletons, reads were clustered into OTUs at 97% similarity and an OTU table was constructed by mapping all labeled reads to these clusters. Taxonomy was assigned to each OTU centroid using the May 2013 version of the Greengenes database and a last common ancestor approach as implemented in QIIME 1.9.1 (McDonald et al 2012). The same pipeline was used to process 18S genes, with a few changes; reads were filtered to a maximum expected error of 0.1 bp per read and no reference-based chimera checking was used. The 18S component of the SILVA database v123 (22-08- 2016) was used for taxonomy assignment (Quast et al 2013).

### Diversity analysis

Downstream analysis was completed in R (Team 2000), using the ‘phyloseq’ (McMurdie and Holmes 2013) and ‘vegan’ packages (Oksanen et al 2013). Samples were rarified to an even depth of 21,600 reads per sample, and 16S rRNA gene amplicons not classified as Bacteria were removed. Counts of unique OTUs and Simpson’s Evenness were used to characterize alpha diversity (Hamady et al 2010). A metric for the successful immigration of taxa was calculated as the fraction of the community composed of OTUs that were previously unobserved on all prior sampling days. This metric is equivalent to taking the finite difference of the sum of observed microbes (i.e., gamma diversity) over time. Beta diversity was measured by calculating unweighted Jaccard distances between samples, and then partitioning those distances into contributes from nestedness (species loss) and turnover (species gain) using the ‘betapart’ R package (Baselga and Orme 2012), followed by the adonis test to measure changes associated with sampling day.

### Confocal microscopy

Confocal fluorescence microscopy was used to confirm the attachment and composition of the biofilm. The slides were stained with 10 μg mL^−1^ Hoechst 33342 (Life Technologies, Carlsbad, CA) for 10 min to target DNA. Images were acquired at 1.5 μm z-steps on a Zeiss LSM 710 scanning head confocal microscope equipped with a Zeiss a Plan-Apochromat 63x/1.40 Oil DIC M27 objective. Excitation lasers were 405 and 633 nm for the blue and red emission channels, respectively. DNA staining was detected at 410-559 nm, and chlorophyll autofluorescence was detected at 647-721 nm. Laser dwell times were 0.64 μs for both channels. Image analysis was conducted using Volocity (PerkinElmer, Waltham, MA).

### Stable isotope probing

Incubations of colonized glass cover slips were performed in sterile HLA-400 medium (Cole et al 2014) with stable isotope tracers were performed with ^13^C labeled HCO_3_- and ^15^N labeled N_2_ gas. A control incubation was established to account for the natural abundance of ^13^C by adding 1.5 ml of a 63.1875 g L^−1^ NaHCO_3_ solution. The labeled incubations were prepared by mixing 0.75 ml of this natural abundance solution with 0.75 ml of a 63.1875 g L^−1^ of ^13^C labeled NaHCO_3_ solution. The ^15^N labeled N_2_ incubations were performed in the HLA-400 medium (minus NH_4_Fe(III) citrate) containing 1 mM NaHCO_3_; 100 ml glass bottles were charged with 20 ml of degassed medium and 1 coverslip sample. A more detailed report of these methods is provided in the supplementary material.

### NanoSIMS imaging

High-resolution secondary ion intensity and isotope ratio maps were generated using a NanoSIMS 50L (CAMECA, Gennevilliers, France) at the Environmental Molecular Sciences Laboratory (EMSL) at Pacific Northwest National Laboratory (PNNL). Biofilm-coated coverslips were mounted on one-inch-diameter aluminum pin stubs and coated with 20 nm of high-purity gold or iridium to improve conductivity. A more detailed report of these methods is provided in the supplementary material.

### Statistics

This study made use of the adonis test (permutation MANOVA) to partition binary Jaccard distance matrices as described above. Generalized linear models were fit to richness, evenness, and within-day βNTI using sampling day as a fixed effect, and using Poisson, gamma, and Gaussian distributions respectively. The ‘multcomp’ package was used to perform *post hoc* multiple comparison testing using Tukey’s HSD (Hothorn et al 2008). We employed a null-model analysis to compare the beta Mean Nearest Taxon Index (βNTI) for all pairwise comparisons within, but not between, the time-points of community development to assess if the phylogenetic similarity between two samples (beta diversity) was significantly higher or lower than expected by chance. The analysis relied on a previously reported convention (Dini-Andreote et al 2015) that reports βNTI values as a z-score with the following interpretation: scores greater than +2 standard deviations from the mean indicate variable selection pressures; scores near zero indicate no significant selection (dominance of stochastic processes); and scores less than −-2 standard deviations indicate homogeneous selective pressures. This method has proven robust in a range of ecosystems (Chase and Myers 2011, Graham et al 2017, Stegen et al 2012). The ‘picante’ package and previously described custom scripts (Stegen et al 2013) were used to calculate βNTI in order to measure changes associated with niche-based processes and selective pressure towards homogenous or heterogeneous phylogenetic structure.

### Data repository and reproducible analyses

Genetic sequencing data is available on the Open Science Framework (osf.io) for both 16S and 18S rRNA gene amplicons as part of this project: https://osf.io/a48vj/. Feature abundance tables of amplicons, along with environmental measurements and scripts used for analysis and graphing are available on GitHub: https://github.com/pnnl/brislawn-2018-founders-species

## RESULTS

### The components and morphology of a developing community

Microbial cells colonized the sterile glass substrate and formed a dynamic, multi-kingdom community over a 79-day succession period. Rich microbial communities were established within 8-days of colonization (Fig. 1A). By this early time point, representatives were identified across 37 bacterial and 16 eukaryotic phyla from over 1013 and 816 bacterial (16S) and eukaryotic (18S) OTUs, respectively. Species richness decreased strongly as the community developed. After a 79-day incubation, the mature microbiome had lost, on average, more than 57% and 39% of the total respective bacterial and eukaryotic OTUs observed from the initial 8-day (colonization) sample. These decreases were statistically significant as determined using *post hoc* general linear hypothesis testing with Tukey’s all-pair comparisons, which yielded p-values < 0.001. The colonizing community was very uneven, as measured by Simpson’s Evenness. This metric ranges between zero and one, with zero representing a community dominated by a single member. The median evenness of 0.02 and 0.03 for 16S and 18S rRNA gene sequences, respectively, emphasizes that a relatively small number of abundant microbes dominated during the development of this community. Bacterial evenness did not mirror the reduction in species richness; evenness only decreased on day-56 (all p-values < 0.001) but rebounded by day-79. No statistically significant changes (all p-values ≥ 0.05) were observed in eukaryotic evenness during the development of this community.

**Figure 1.**
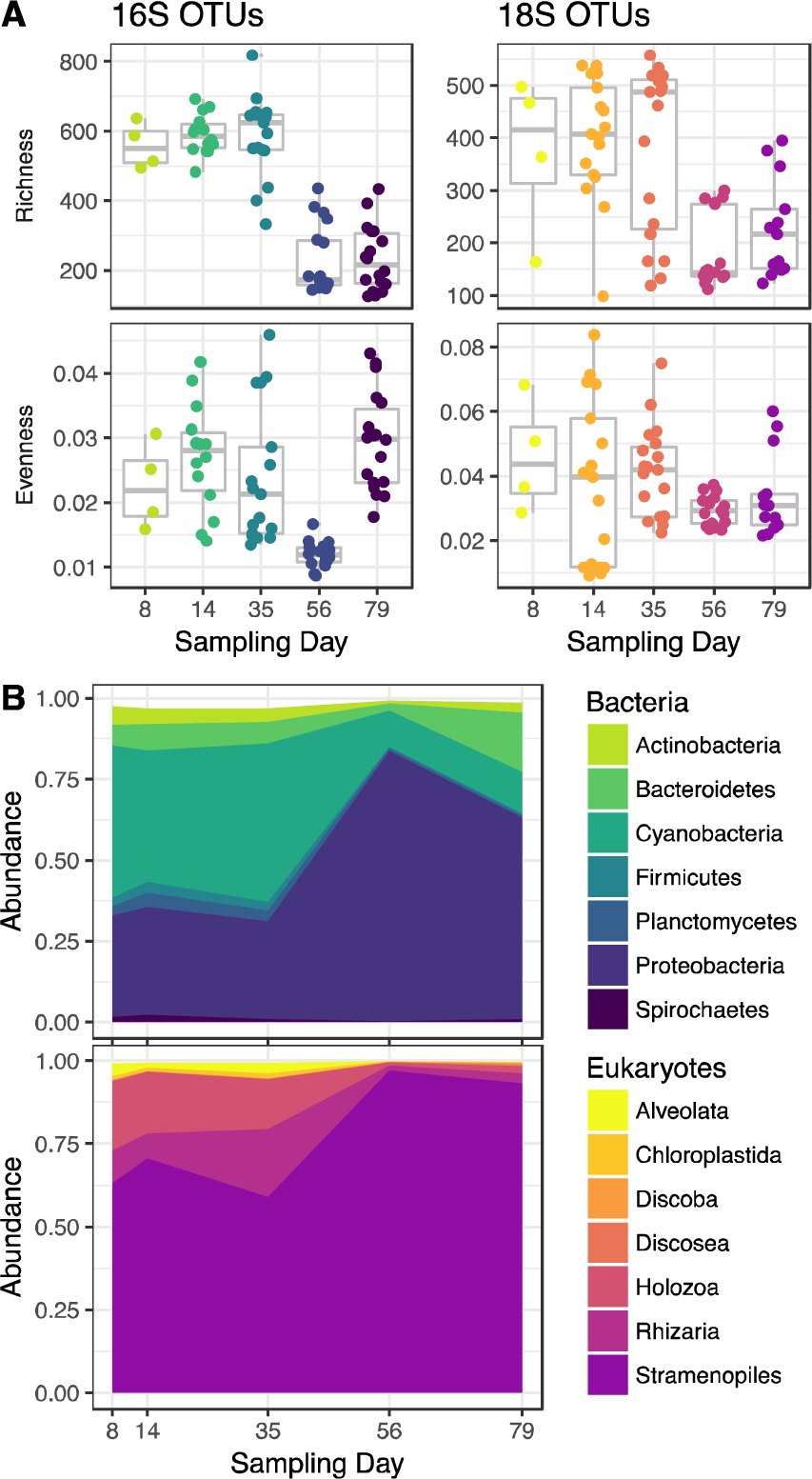
The bacterial and eukaryotic composition of the biofilm community changed during the observed succession period. **A)** Changes in alpha diversity as described by species richness (unique OTUs) and Simpson’s evenness. **B)** Changes in the relative abundance of common bacterial and eukaryotic taxa identified from 16S and 18S OTUs.

Cyanobacteria and Proteobacteria were the dominate bacterial phyla over the observed maturation period (Fig. 1B). Some of the most common bacteria had greater relative abundances during the early (colonizing) time periods but then decreased with time. These included: Actinobacteria, Firmicutes, Planctomycetes, and Spirochaetes. The most abundant eukaryotes, through all phases of community development, belonged to the superphylum Stramenopiles. Interestingly, this broad phylogenetic group increased its relative abundance as the microbiome matured (Fig. 1B), while the remaining eukaryotic taxa decreased in relative abundance.

The morphology and spatial distribution patterns of inorganic carbon and N_2_ incorporation also changed during the 79-day succession. Confocal micrographs showed that diatom-like cell structures became more frequent as the community matured (Fig. 2A). Interestingly, a shift in morphology was observed; larger diatom-like cells common to the 14- and 35-day samples were replaced by a more dominant and smaller diatom-like cell structure in the final sampling points. This imaging based result corresponded to the molecular evidence for the apparent takeover of Stramenopiles in the latest stages of observed succession (Fig. 1B). The magnitude and distribution of ^13^C and ^15^N incorporation was measured by using ^13^C-labelled bicarbonate and ^15^N-labelled N_2_ tracers in combination with nano-SIMS imaging (Fig. 2B and supplementary file). These results show that autotrophy becomes more prevalent throughout the course of community development and corresponds with higher occurrence of diatom-like structures. Primary producers were not prerequisite for the recruitment of new cells to the biofilm. Relatively frequent occurrences of N_2_ fixation was observed during all phases of community development indicating that at least a subset of the founding species was diazotrophic.

**Figure 2.**
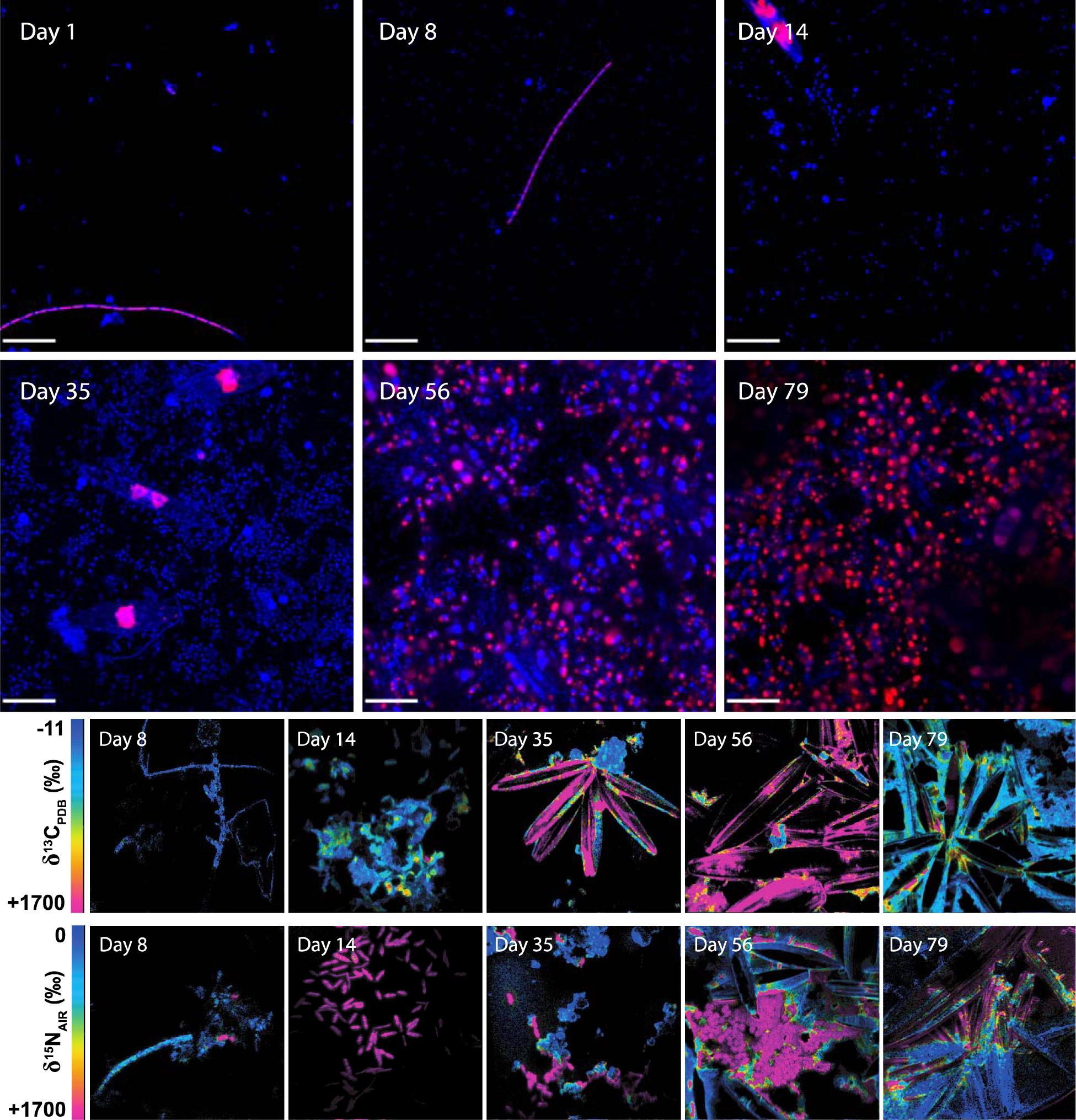
The community changed morphology frequency of autotorphic and diazotrophic cells as members were recruited during succession. **A)** Confocal micrographs showing changes in biofilm community morphology as visualized by the DNA stain (Hoechst; blue) and chlorophyll autofluorescence (red); scale bars represent 20 µm. **B)** Magnitude and distribution of ^13^C and ^15^N incorporation in representative biofilm communities showing incorporation of labelled substrates over different incubation periods. Top row: δ^13^C distribution after incubation with ^13^C-labelled bicarbonate. Bottom row: δ^15^N distribution after incubation with ^15^N-labelled N_2_. The full suite of replicate nanoSIMS images are presented in a supplementary file.

### Which ecological processes drive community structure changes?

Changes in the community over the maturation period were attributed to multiple processes including nestedness, and turnover. Decreases in species richness observed over the community’s maturation inspired us to ask how successful immigration changed over time. In this context, successful immigration is governed by dispersal of regional taxa, followed by attachment and growth. Dispersal is the movement of microbes across space, specifically, onto the glass slides from the surrounding environment. We answered this question by employing a relatively simple inference of successful immigration, which counts the fraction of the community represented by OTUs that were undetected at any previous time point (Fig. 3A). This analysis showed that the recruitment of new OTUs diminished 50% by day-14. The successful immigration of bacteria and eukaryotic species continued to decrease during the maturation period and was negligible by day-56 (only 3.3% of the observed OTUs were new). One limitation to this result is the underlying assumption that the appearance of new OTUs was actual due to dispersal and not caused by blooms of taxa residing below detection.

**Figure 3.**
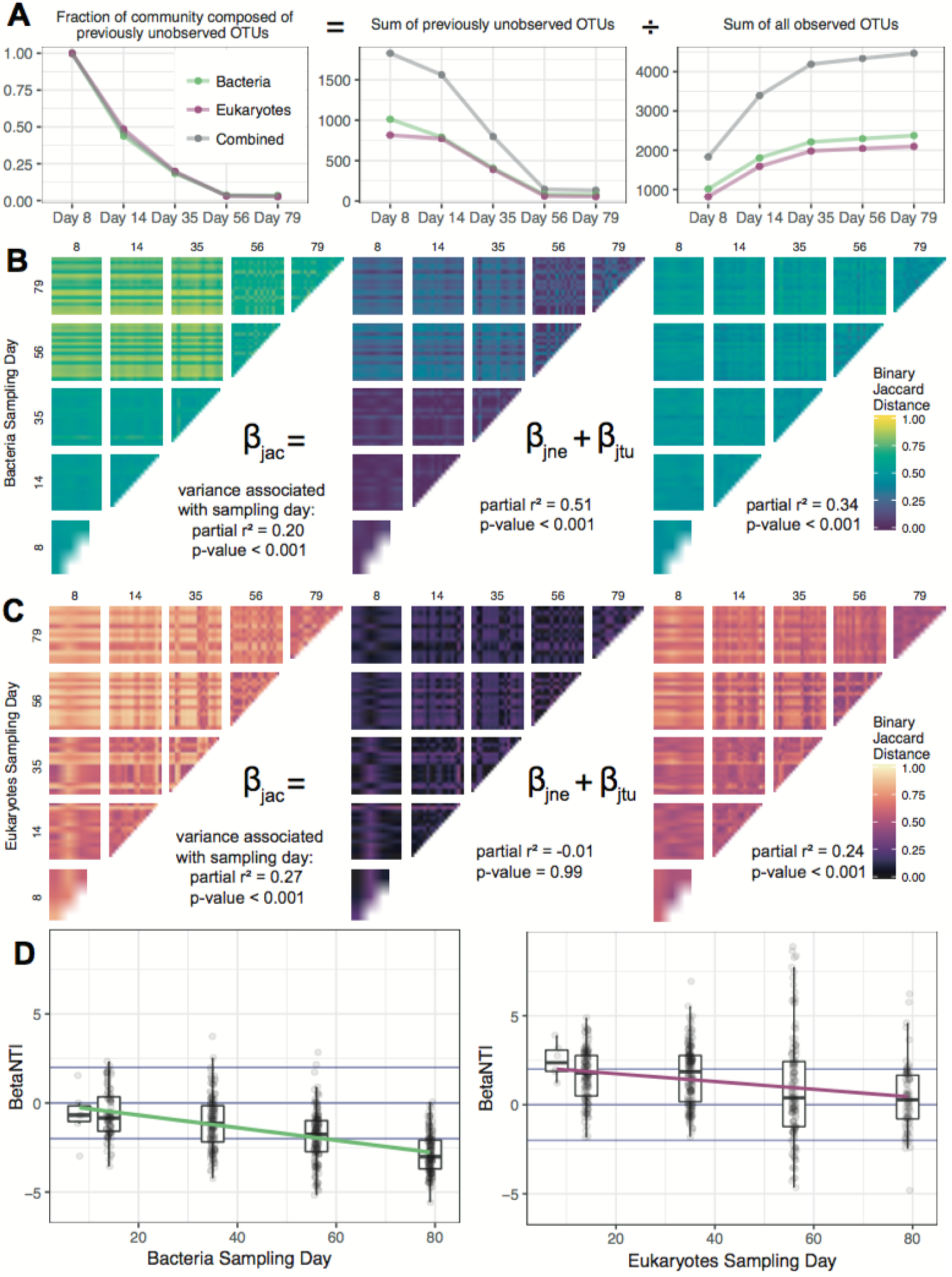
Different ecological processes contributed to the observed beta diversity depending on the stage of community succession and faction of the microbiome (bacteria or eukarya). **A)** The successful immigration of taxa over time as estimated by introduction of new OTUs. **B)** Binary Jaccard distance between samples, partitioned by contributions from nestedness (species loss) and turnover (species replacement). **B)** Inferred changes in degree to which variable selection (βNTI > + 2), stochastic processes (−2 < βNTI < +2) and homogenous selection (βNTI < −2) contribute to different factions of the community (bacteria and eukarya) at each observed stage of succession.

While the patterns of new OTU recruitment are informative, they do not directly account for how much temporal variation in community structure was due to the gain and/or loss of OTUs. For this, we employed a more rigorous analysis of beta diversity by accounting for the additive contributions of nestedness (species loss) and turnover (species replacement) to the binary Jaccard distance (total beta diversity) calculated between all samples taken at all time points (Baselga 2012). The adonis test was used to compare the amount of variation from total Jaccard distance, nestedness, and turnover that could be attributed to sampling day (Oksanen et al 2013). Turnover was a much greater contributor to the observed beta diversity than nestedness with respect to both the bacterial and eukaryotic components of the community. However, within the bacterial samples, more than 50% of the variance observed from nestedness could be attributed to sampling day (Fig. 3). In comparison, only 20% of turnover and 34% of total Jaccard could be attributed to differences observed between sampling days. Hence, while bacterial turnover was high, it was relatively constant between each successional period. Bacterial nestedness was more variable between samples and changes to bacteria community structure coincided more with changes to nestedness. Specifically, there was an increase during the drop in alpha diversity on day-56. We observed the opposite result with respect to changes in eukaryotic beta diversity assessed between sampling days, which was attributed more to turnover. Eukaryotic nestedness was essentially constant and showed little association with time (Fig. 3BC). Observations from the bacterial components of the microbiome were consistent with the temporal diminishment of successful immigration and showed that the structure of the community was driven by the OTUs gained in the early phases of development.

We were curious as to whether bacterial and eukaryotic succession were influenced more by environmental selection or stochastic processes. We know that if we assume dispersal to be a stochastic process – which is a reasonable assumption for this benthic ecosystem – then the earliest stages of succession were at least partially influenced by stochastic processes. However, this insight does not tell us if stochastic processes such as dispersal outweighed environmental filtering. If the founder species exerted strong homogenous selection pressures onto their environment, then taxa recruited in the subsequent stages of succession would be expected to be more phylogenetically related than would occur by chance (Andersson et al 2010, Stegen et al 2012). In contrast, if stochastic process dominated, then the effect of abundant colonizers would be damped such that no phylogenetic pattern in successive OTU recruitment would be expected. By measuring the phylogenetic similarity of our bacterial and eukaryotic OTUs, we were able to infer the strength of selective pressure and distinguish the relative contribution of stochastic versus deterministic processes during the development of this community (Dini-Andreote et al 2015, Graham et al 2017). Stochastic processes were found to dominate bacterial colonization (−2 < βNTI < −2 at early time points) but trended towards deterministic, homogenous selection (βNTI < −2) in the latest stages of succession (Fig. 3D). The eukaryotes appeared to be influenced by an opposite temporal ordering of stochastic and deterministic process during the observed succession period. The within time-point βNTI values showed that eukaryotic colonization processes were initially influenced by weak variable selection (βNTI > +2) with a significant mixture of stochastic and deterministic processes at work as the community matured (Fig. 3C).

### Who were the founder species and how well did they persist?

We determined which taxa had the highest relative abundances (> 2%) from the initial period of colonization defined by the 8-day sampling point. As mentioned, these abundant colonizers were defined as the founder species (Fig. 4). An identical analysis was performed on the 79-day samples. This threshold identified only four and five bacterial and eukaryotic taxa ranked at the Family level. Of these nine Families, only three maintained high relative through the observed succession period. As the community matured, the remaining majority of the founder species (six of nine) were replaced by taxonomic families that were found only at low relative abundance at the 8-day colonization sampling point. The most abundant taxa observed in the community did not bloom until the 56-day sampling point indicating a clear pattern of succession (Fig. 4 CD).

**Figure 4.**
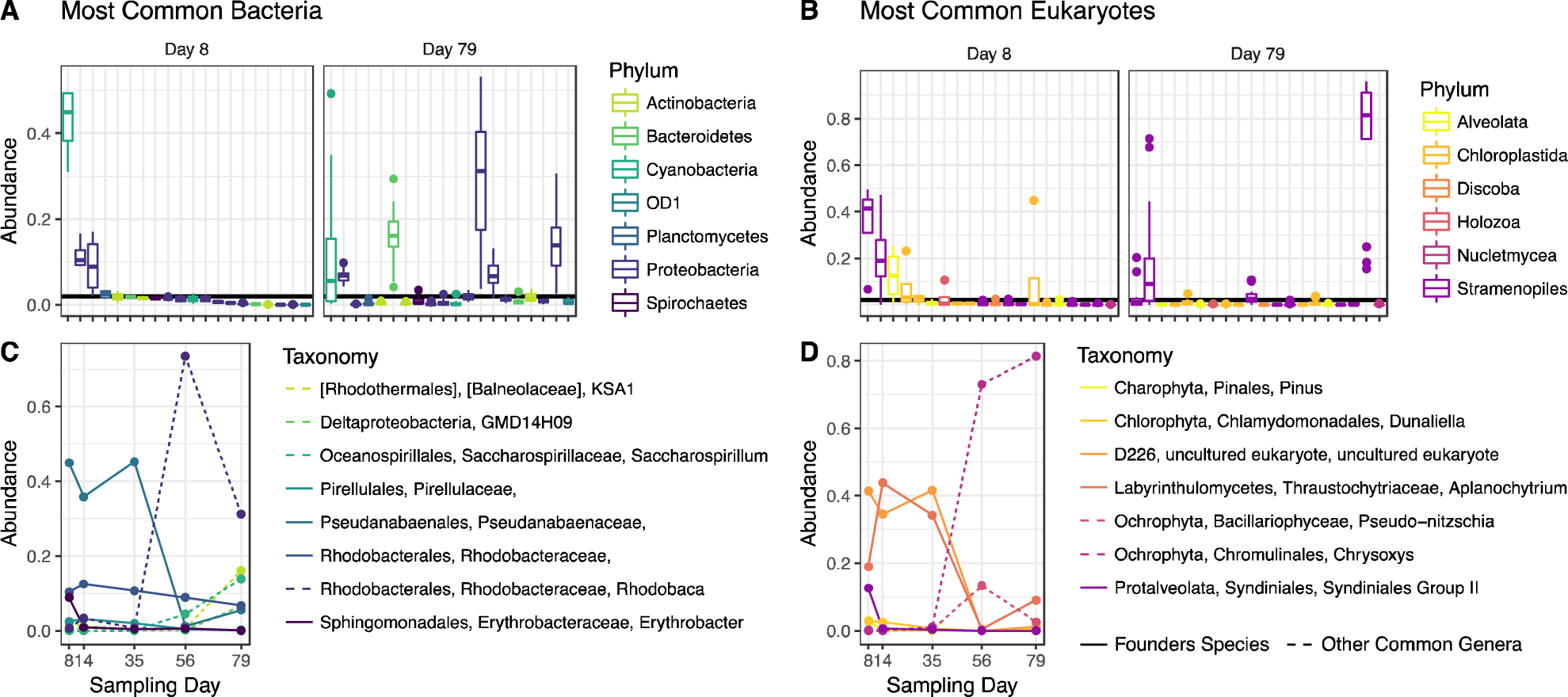
Identification of founder species and their abundances during maturation. **A and B)** Ranked abundances of common bacterial and eukaryotic families, color-coded by phyla. A threshold relative abundance value was set at 2% and the most abundant OTUs belonging to taxa that exceed the threshold by day-8, which was chosen as the representative time for colonization, are defined as founder species. The persistence of founder species is compared to the relative abundances of other taxa that were able to exceed the threshold by day-79, the end point for this study. **C and D)** Temporal changes in abundances of specific founder species (solid lines) compared to other highly abundant OTUs that emerge over the microbiome’s maturation.

The most abundant founder species were annotated as Pseudanabaenaceae for bacteria and two Stramenopiles (Aplanochytrium and an uncultured D226) for eukaryotes. Interestingly, they maintained high relative abundances for the first 2-4 weeks, but were ultimately replaced by Rhodobacteraceae, Rhodobaca and Chromulinales, Chrysoxys, which at some points comprised more than 70% of their respective kingdoms. All founder species showed similar dynamics; initial success (in terms of relative abundance) followed by replacement. Yet, this replacement was not absolute; both Pseudoanabaenaceae and the Stramenopiles remained dominant in some samples over the observed succession period. This persistence throughout months of consistently high turnover (Fig. 3B) demonstrates their ability to invoke a founder’s effect; even as the selective pressure towards a homogenous community increased (Fig. 3D) Pseudanabaenaceae was the lone bacterial founder species able to persist at high relative abundance. Interesting, the type of selection differs between kingdoms. Initially, bacterial founder species experience a minimal amount of phylogenetic selective pressure, and selection towards related microbes slowly increases (Supplemental Fig. S6 and Table S2). Eukaryotes experience a different selective pressure; the Chrysoxys family of algae along with the diatom Pseudo-nitzchia became dominant in the later phases of community development, when selection based on niche is minimal and stochastic processes dominated (Supplemental Fig. S7 and Table S3).

## DISCUSSION

This study was designed to ask if the early abundance of colonizers (founder species) relate to persistence or turnover. We found that six of nine identified founder species did not maintain high relative abundances during the observed succession. Hence most of the abundant colonizers forfeited their founder’s effect. We also investigated the degree to which deterministic and/or stochastic processes dictate the roles that founder species play in their mature communities. We postulated that if a founder species was able to influence community succession by imposing strong selection, then we would see a greater degree of phylogenetic relatedness of new taxa recruited in the next stages of community succession. The null models indicated that colonization of the bacteria was likely not influenced by strong selection but rather by stochastic processes. Likely the initial selective forces were overwhelmed by dispersal and a high degree of taxonomic sampling from the contiguous benthic habitat and sedimentation from overlaying water column. However, the null models indicated that successive stages in the bacterial community development trended towards homogenous selection. Hence, the bacterial founder species likely modified their localized environment in ways that mediated environmental filtering – exerted priority effects. Homogenous selection appeared to act more strongly on the bacterial components of the community in later stages of succession when the biofilm community had presumably sampled the majority of taxa from the nearby environment. We note that founder species – as well as other abundant taxa identified in this study – were identical or highly related to some of the most common taxa reported by previous studies performed in the Hot Lake ecosystem in different years (Bernstein et al 2017a, Lindemann et al 2013, Mobberley et al 2017, Moran et al 2014). This indicates that, although the phyla are shown to decrease during this 79-day succession period, these taxa are likely stable and persistent within the ecosystem.

Interestingly, the ecological forces acting on bacterial succession were found to be in sharp contrast to the results obtained for the eukaryotic colonizers. We concluded that weak variable selection forces were initially imposed onto the eukaryotes followed by a mixture of stochastic and deterministic processes during later stages of succession. One possibility that must be considered is that by day-79, bacteria ended the known succession period that was linked to previously identified seasonal assembly (Lindemann et al 2013), but that the recruitment and succession of eukaryotes operated on different time scales. This would explain why the bacterial and eukaryotic βNTI curves exhibited negative slopes indicating a trend towards convergent homogenous selective pressures through time. This also implies that the dominant eukaryotic blooms occur after bacterial recruitment had stabilized. Perhaps bacterial taxa found in high abundance in the later stages of succession established the niches required to recruit diatoms and other abundant eukaryotes. The microscopy provided qualitative evidence of different kingdom-specific time scales as diatom-like structures became common only in the later weeks of the season.

Differences in the relative contribution of turnover and nestedness – attributed to inter-sample diversity – also supported the idea that eukaryotic succession not only operates on a different time scale than the bacteria but also by different sets of ecological processes. We note that the eukaryotic components of the Hot Lake microbiome have not been characterized as thoroughly as the bacteria. However, the abundant eukarya identified here were consistent with one other study of the Hot Lake ecosystem, performed at a similar sampling location and on a different year (Bernstein et al 2017b). Another important consideration for our interpretation of turnover and nestedness is the fact that these measurements (as applied here) only accounted for presence/absence of taxa. We know that the majority of OTUs are found in low abundance, but we do not know how many taxa are undiscovered and below detection limits of our amplicon analyses. The low abundance and undiscovered taxa can serve as a seed bank, blooming to replace others as conditions change (Shoemaker and Lennon 2018). Those that may have escaped detection could create artifacts in our interpretations. However, within the limitations of our current measurement technologies we are able to determine that turnover and nestedness are important to microbial community succession dynamics and drive shifts in taxonomic abundance.

Alpha diversity decreased as the community matured. This was shown by decreases in species richness and the diminishing rate of new OTU recruitment. Similar observations have been made in other aquatic biofilm succession studies (Besemer et al 2007, Lyautey et al 2005) that also started from a bare substratum. This reoccurring observation presents the possibility of a broader principle in that the conditions that favor recruitment in biofilm rich ecosystems select for those taxa that have the ability to adhere to surfaces (Niederdorfer et al 2016). It also expected that that new niches are created during the initial stages of biofilm community succession as these surfaces are filled (Niederdorfer et al 2017). The biofilms observed from this current study exhibited clear distinctions between the succession patterns of bacteria and eukaryotes. This was observed from both molecular and imaging evidence. Diatoms were recruited during the mid-development phase after bacteria-like cell structures had attached. Unfortunately, these current results are not sufficient for determining inter-kingdom relationships; however, we expect that they are coupled and that they influence each other’s recruitment to the biofilm.

Algal taxa were by far the most abundant eukaryotes identified, yet the microscopy analyses did not reveal diatom-like structures to be common until the mid-development stages of this community (beginning on day-35). The images also qualitatively show uptake of ^13^C-label increases with the arrival of frequent diatom-like cell structures. While it is not surprising that microalgae act as primary producers, the finding does adds context to this unique hypersaline ecosystem where previous reports have mostly attributed primary production by cyanobacteria (Bernstein et al 2017a, Lindemann et al 2013, Moran et al 2014); but see (Bernstein et al 2017b). The importance of microalgae to the ecosystem is also supported by our identification of founder species, including photosynthetic eukaryotes (Charophyta or Pseudo-nitzchia) make up at least 40% of 18S OTUs in any given sample. This is not to imply that autotrophic bacteria do not play a role in primary productivity, as 16S OTUs belonging to the Pseudoanabaenaceae were highly abundant in the current study and previous reports from the Hot Lake ecosystem (Bernstein et al 2017a, Bernstein et al 2017b). It seems plausible that the diazotrophy could be more important than autotrophy as a pioneer function for creating new niches required for species recruitment; many of the abundant cyanobacteria, including those belonging to Pseudanabaenales, are either taxonomically related to known nitrogen fixers or implicated by a previous metagenomics study (Mobberley et al 2017, Stal 2009). A precedent for this idea has been described by microbial succession studies in other ecosystems from relatively extreme environments including acid mine drainage and glacial forelands (Brankatschk et al 2011, Duc et al 2009, Huang et al 2011). This concept is partially supported by the eukaryote-like cell structures were found to be in close physical proximity to diazotrophic cells signifying tightly associated food webs with respect to multi-kingdom nitrogen cycling.

Another goal of this study was to elucidate the ecological processes motivating changes in structure and basic function of the microbiome from colonization to maturation over the observed succession period. The basic functions interrogated were limited to autotrophy and nitrogen fixation based on previous research that indicated that potentially nitrogen fixing, filamentous cyanobacteria dominated the Hot Lake microbial mat (Lindemann et al 2013, Mobberley et al 2017). To capture this, we have assembled a narrative to comparatively interpret which ecological processes underpin community development by dividing the community maturation period into three distinct stages: colonization (defined by the 8-day sample), development (defined between the 14- and 56-day samples) and mature (defined by the 79-day sample). The colonization stage was characterized by the arrival of bacteria-like cell structures. Many of which were found to fix N_2_ and were not exclusively autotrophic as shown by the isotope tracer incubations and nanoSIMS images. At least some of the colonizing species were heterotrophic which indicates that primary producers are not necessarily required for recruitment of other cells. This observation is somewhat contrary to other, similar succession studies performed *in situ* (Beam et al 2016, Roeselers et al 2007) but is not surprising based on our knowledge of high abundances of dissolved organic carbon in Hot Lake (Anderson 1958, Zachara et al 2016). The development stage was characterized by the arrival and increased abundance of diatom-like cell structures a diminished rate of successful immigration of taxa from the surrounding environment. The maturation stage was characterized by the effective loss or turnover of most founder species and a contrasting shift in the stochastic versus deterministic ecological processes as compared to colonization; specifically, bacteria shifted from stochastic towards homogenous selective and eukarya shifted from weak-variable selection towards stochastic processes.

## Conclusions

The identity and roles of the most abundant colonizers in microbial communities – i.e., founder species – are often overlooked. In this succession study, we found that both the bacterial and eukaryotic founder species were mostly turned over. We also found that that each kingdom appeared to be influenced by different ecological forces and time scales of succession. Specifically, bacteria shifted from stochastic towards homogenous selective and eukarya shifted from weak-variable selection towards stochastic processes. Successive stages in the bacterial community development trended towards homogenous selection. Hence, the bacterial founder species likely modified their localized environment in ways that mediated environmental filtering where eukaryotic founder species may not have played as great of a role in the later stages of succession.

## ACKNOWLEDGMENTS

The authors would like to thank Joe Brown and James Fredrickson for valuable discussions and technical assistance. The authors would also like to acknowledge the U.S. Bureau of Land Management, Wenatchee Field Office, for their assistance in authorizing this research and providing access to the Hot Lake Research Natural Area.

