## Supplementary Materials for "Forfeiting the founder effect: turnover defines biofilm community succession" (from another version of this preprint: version 1, 2018-03-15)

**Supplementary Data and Methods for:** Forfeiting the founder effect: turnover defines biofilm community succession

^1^Biological Sciences Division, Pacific Northwest National Laboratory, Richland, WA, USA; ^2^Signature Science and Technology Division; Pacific Northwest National Laboratory, Richland, WA, USA; ^3^ Environmental Molecular Sciences Laboratory, Pacific Northwest National Laboratory, Richland, Washington, USA; ^4^The Gene and Linda Voiland School of Chemical Engineering and Bioengineering, Washington State University, Pullman, WA, USA;

**** Correspondence****:* Hans C. Bernstein, Biological Sciences Division, Pacific Northwest National Laboratory, P.O. Box 999, MS-IN: J4-18, Richland, WA 99352; James J. Moran, Chemical and Biological Signature Science, Pacific Northwest National Laboratory, P.O. Box 999, MS-IN: P7-50, Richland, WA 99352

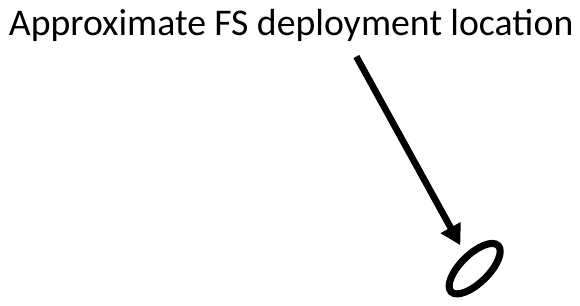
**Figure S1.** The approximate location of in situ incubations place within Hot Lake (north-central Washington State, USA; 48.973062°N, 119.476876°W). Incubations were performed on sterile, borosilicate surfaces deployed in 43-50 cm water depth.

**Detailed methods for stable isotope probing.** Petri dish containing glass coverslips were harvested from each time point and transferred (in 4°C coolers) to a mobile laboratory within 2-6 h of sampling. Incubations with stable isotope tracers were performed with ^13^C labeled HCO_3_- and ^15^N labeled N_2_ gas. The glass coverslips sampled from the lake at each time point were divided between two 950 ml sterile pyrex dishes, each charged with 500 ml of a previously described HLA-400 (Cole et al 2014) medium minus NaHCO_3_, TES buffer and NH_4_Fe(III) citrate. A control incubation was established to account for the natural abundance of ^13^C by adding 1.5 ml of a 63.1875 g L^-1^ NaHCO_3_ solution. The labeled incubations were prepared by mixing 0.75 ml of this natural abundance solution with 0.75 ml of a 63.1875 g L^-1^ of ^13^C labeled NaHCO_3_ solution (lot number Pr-24217, Cambridge Isotope Laboratory, Tewksbury, MA). The ^15^N labeled N_2_ incubations were performed in the HLA-400 medium (minus TES and NH_4_Fe(III) citrate) containing 1 mM NaHCO_3_; 100 ml glass bottles were charged with 20 ml of degassed medium and 1 coverslip sample. Each bottle was capped with a septum and purged with N_2_ for 10 min; the head space was charged with 20 ml (1 ATM) of ^15^N_2_ (98 atom % 15N, product number 364584, Lot: MBBB0968V, Sigma-Aldrich, St. Louis, MO) using a needle and gas-tight syringe. Natural abundance controls were prepared identically by omitting the 20 ml injection of ^15^N_2_ labels. The N_2_ incubation bottles were wrapped in aluminum foil to maintain darkness and anoxic conditions. All incubations were performed for 6 h under a T5 grow light (40 µmol photons PAR m^-2^ s^-1^; Hydrofarm, Inc., Petaluma, CA, USA). Following the incubation, the NaHCO_3_ incubated coverslips were removed using sterile forceps and rinsed in 75 ml tissue culture dishes primed with the HLA-400 medium for 30 min to wash away unincorporated bicarbonate; rinse solution was removed. At the conclusion of all incubations and rinses each coverslip was placed in a 3.5 ml of fresh medium containing 4% paraformaldehyde and stored at 4°C.

**Detailed methods for NanoSIMS imaging.** High-resolution secondary ion intensity and isotope ratio maps were generated using a NanoSIMS 50L (CAMECA, Gennevilliers, France) at the Environmental Molecular Sciences Laboratory (EMSL) at Pacific Northwest National Laboratory (PNNL). Biofilm-coated coverslips were mounted on one-inch-diameter aluminum pin stubs and coated with 20 nm of high-purity gold or iridium to improve conductivity. Because of widely varying conductivities of the individual samples several different strategies were employed to maximize secondary signal. Samples with acceptable conductivity were presputtered with 5 × 10^15^ ions cm^-2^, before 10 40 × 40 μm planes were acquired with a dwell time of 2.7 ms pixel^-1^ and at a density of 256 × 256 pixels. Samples from day-8 and day-79 samples were very insulating and a different analysis strategy was employed. These samples were presputtered with 1 × 10^16^ ions cm^-2^, followed by 2 planes of analysis employing 13.5 ms pixel^-1^ dwell time, with a normal incidence electron gun employed for charge neutralization An in-house yeast standard ( δ^13^C = -11.0‰, δ^15^N = +0.35‰) was analyzed daily using conditions identical to the analysis of the biofilms to correct for instrumental mass fractionation. Analysis conditions included a 16 keV, 2 pA Cs^+^ primary ion beam with ~120 nm diameter, a D1 aperture of 200 μm, a 30 μm entrance slit, 200 μm aperture slit, 100 μm exit slits and a centered energy window of ~6 eV (FWHM). Electron multiplier (EM) detectors measured intensities of ^12^C_2_^-^, ^12^C^13^C^-^, ^12^C^14^N^-^, ^12^C^15^N^-^, ^28^Si^-^, ^31^P^-^, and ^32^S^-^ secondary ions in multicollection mode with NMR magnetic field regulation employed. Particular care was taken to resolve ^12^C^15^N^-^ from ^13^C^14^N^-^, ^12^C^13^C^-^ from ^12^C_2_H^-^, and P^-^ from ^30^SiH^-^. m/z scans were performed frequently during each analytical session to assure that instrumental drift did not compromise data integrity. Image processing and analysis were conducted in ImageJ using the OpenMIMS plugin, available from the National Resource for Imaging Mass Spectrometry (NRIMS, http://www.nrims.harvard.edu). All images were corrected for deadtime and quasi-simultaneous arrival (QSA) effects (β = 0.5) and stack-aligned using a best fit alignment algorithm within the plugin. Isotope ratio distributions were converted to Hue-Saturation-Intensity (HSI) image maps generated from the ratios of masses ^12^C^13^C^-^ /^12^C_2_^-^ and ^12^C^15^N^-^ /^12^C^14^N^-^ for carbon and nitrogen isotopes, respectively. Isotope values of individual cells were extracted from pixel-averaged Regions of Interest (ROIs) manually drawn around corresponding image areas and calibrated to cell-averaged yeast standard values obtained from the standard yeast image for that session. Isotope ratios are expressed in traditional delta notation relative to the Vienna Pee Dee Belemnite (VPDB) and atmospheric N_2_ (AIR) international reference standards for carbon and nitrogen, respectively.

**Environmental measurements collected from field.** We took measurements of the lake and mat environment at every time point corresponding to the molecular measurements presented in the main text. The pH, temperature, conductivity, and incident irradiance was also measured using a Sond (MS5, Hydrolab, Loveland, CO,) and a radiometer (LI-COR 250-A light meter and Quantum sensor, LI-COR Biosciences, Lincoln, NE) in the lake water overlaying the sampling position at each time point.

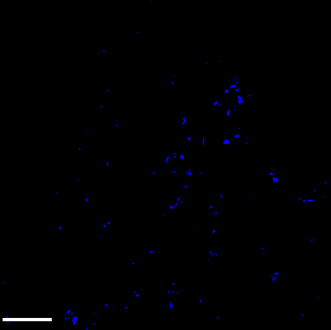

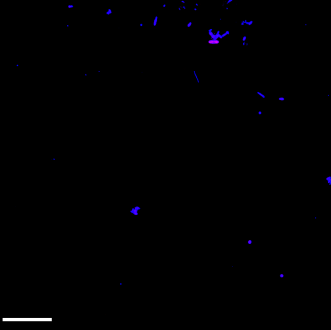

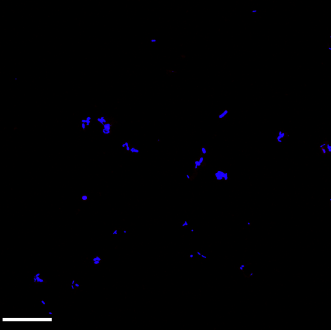

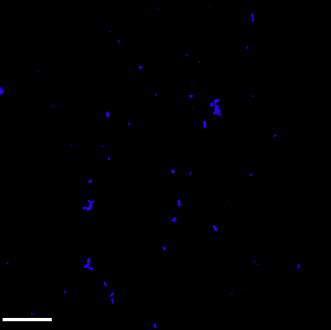

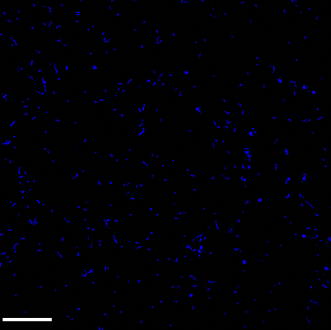

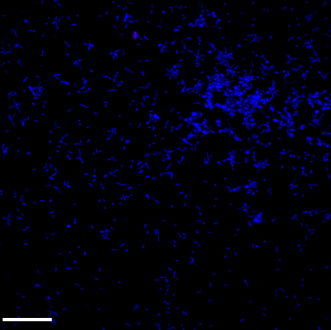

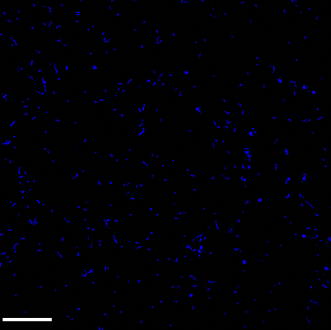

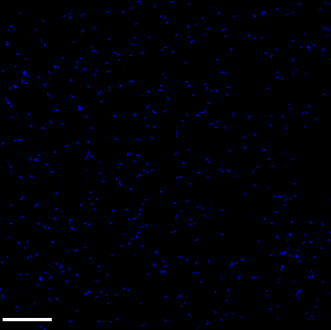

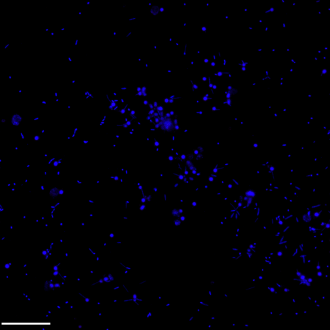

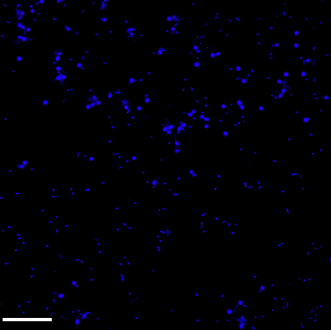

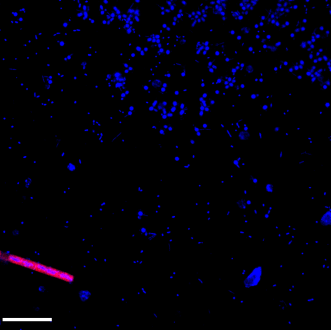

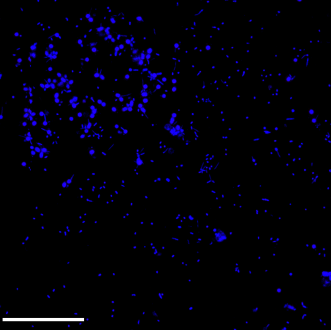

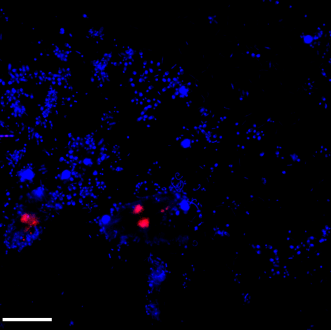

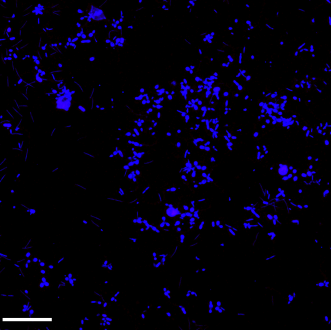

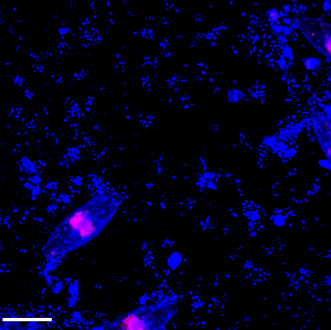

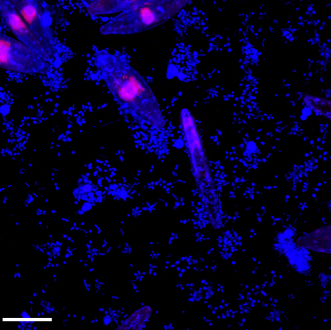

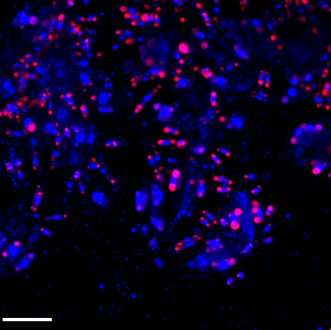

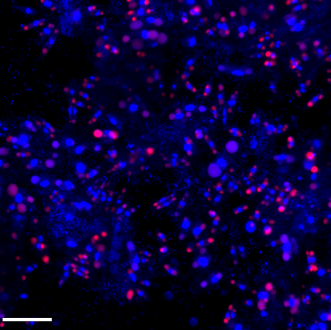

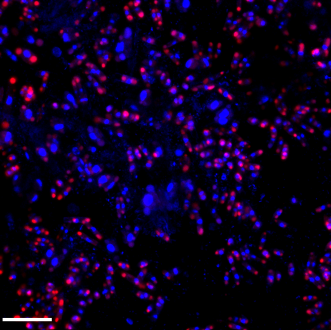

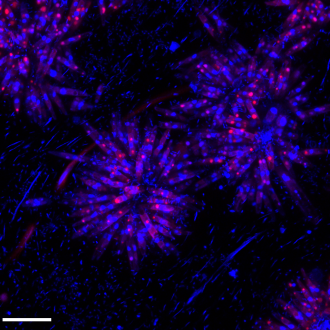

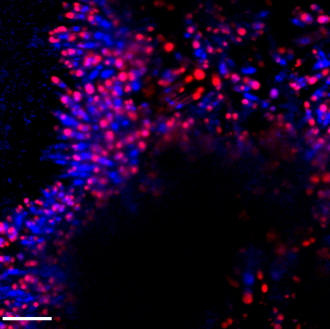

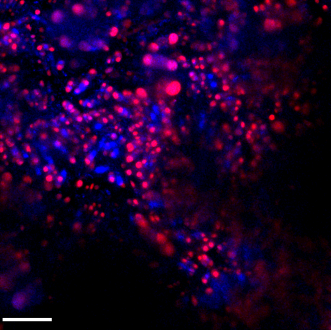

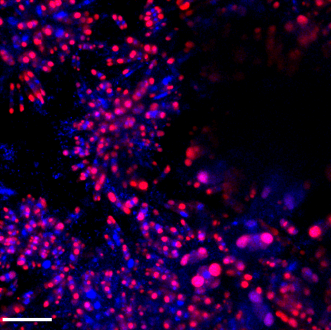

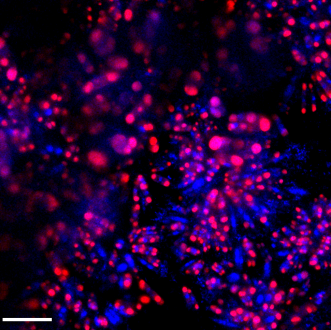

**Figure S2.** The comprehensive collection of confocal micrographs showing changes in biofilm community morphology as visualized by the DNA stain (Hoechst; blue) and chlorophyll autofluorescence (red); scale bars represent 20 µm. The rows of images or organized in descending order and correspond to the following experimental sampling points: 1-day, 8-day, 14-day, 35-day, 56-day and 79-day samples. Each image is 1 of 5 replicate frames; four are shown here and the remainder are shown in within Figure 2 of the main text.

**A) ^13^C-labeled bicarbonate incubation**

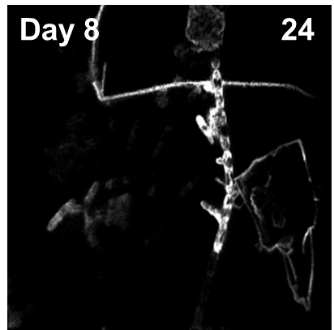

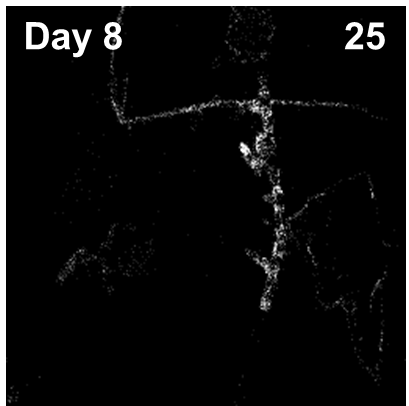

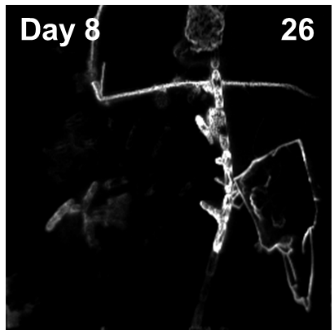

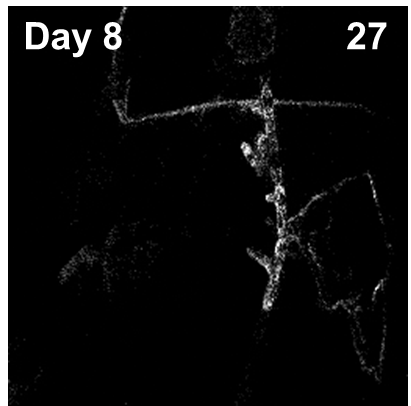

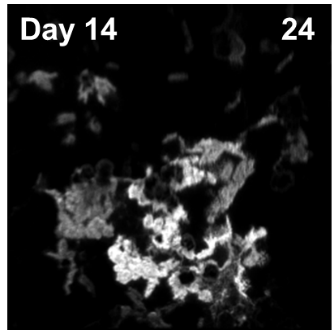

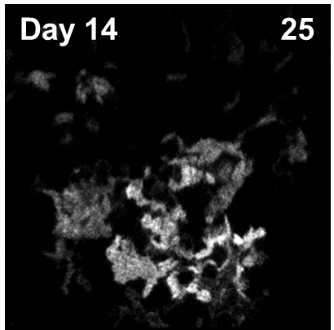

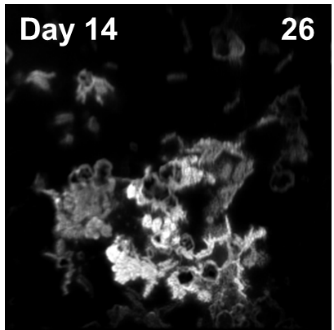

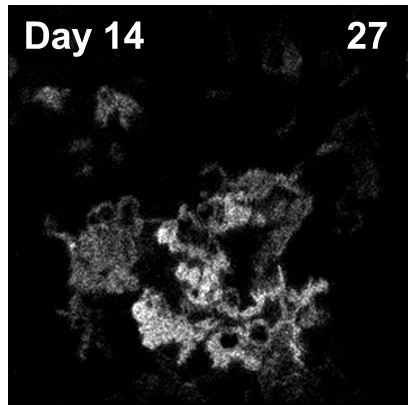

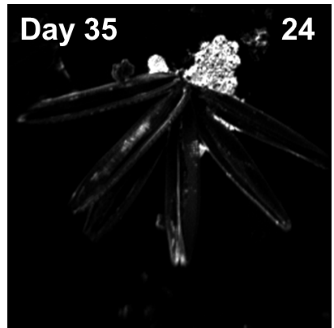

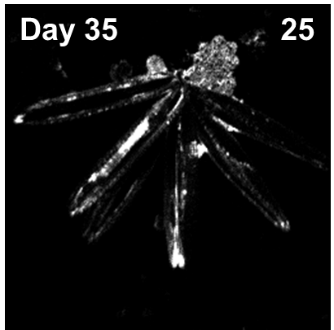

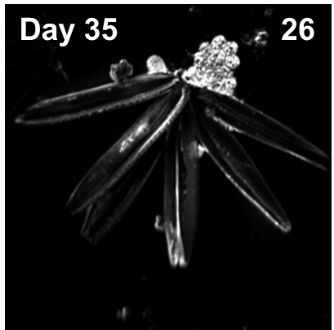

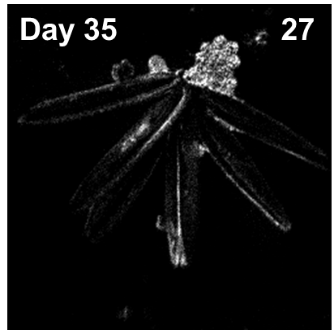

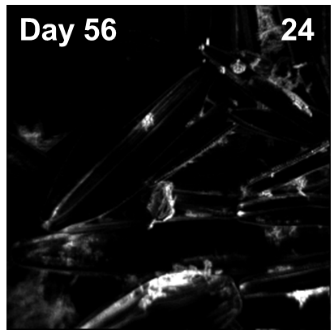

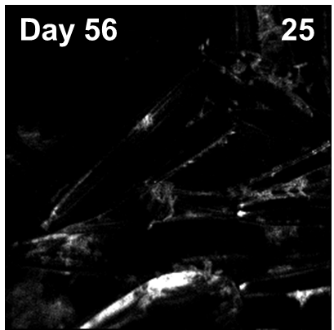

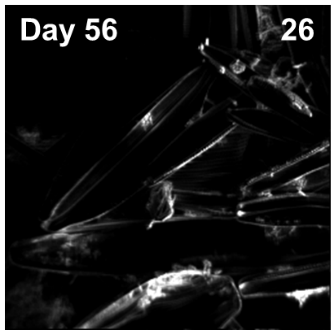

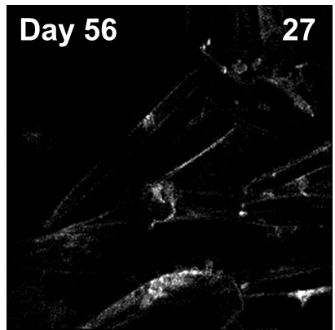

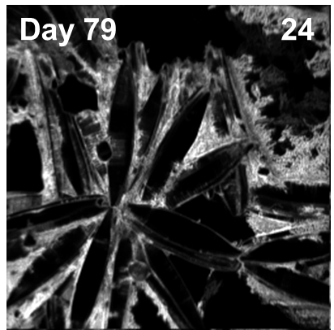

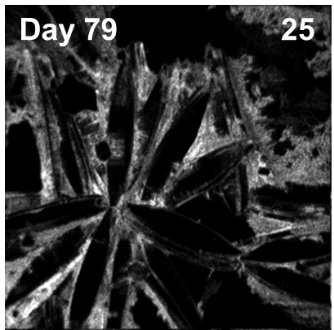

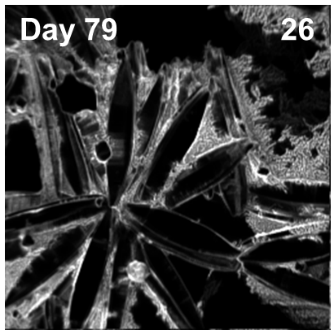

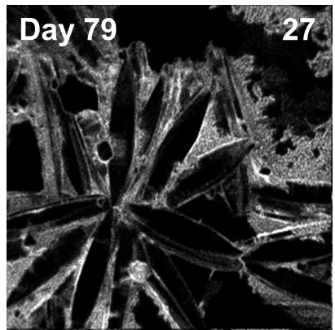

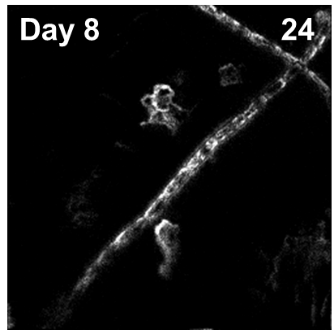

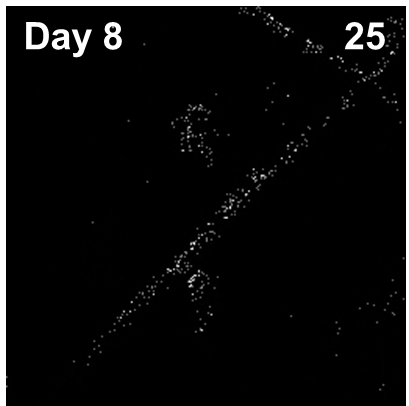

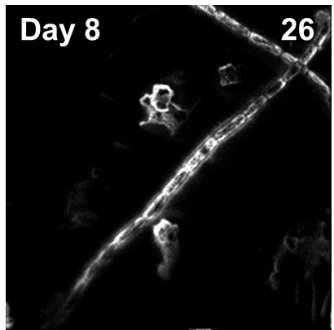

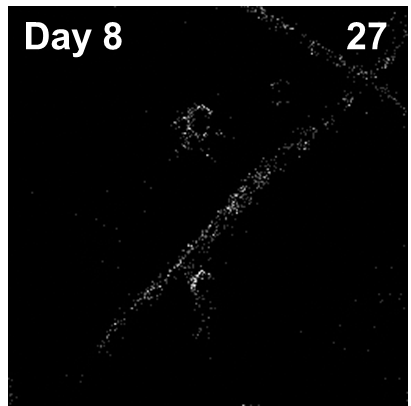

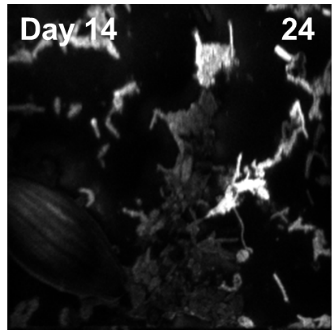

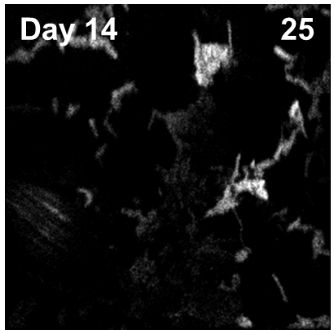

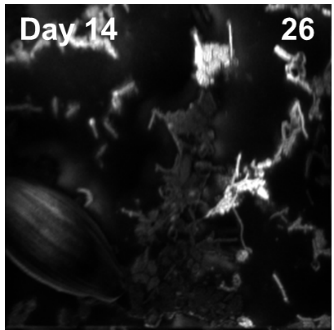

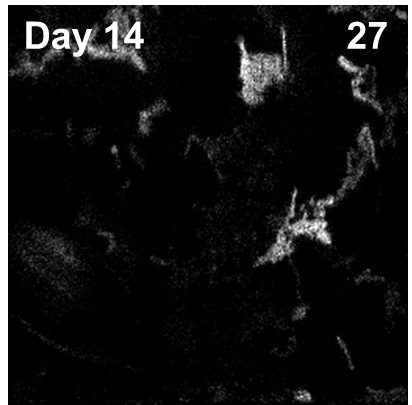

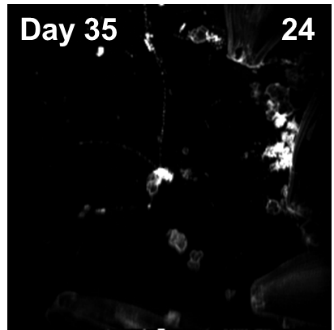

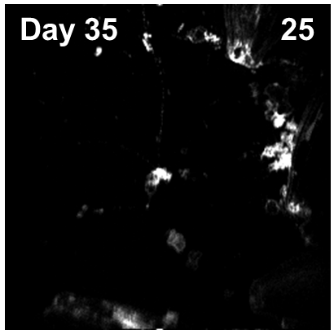

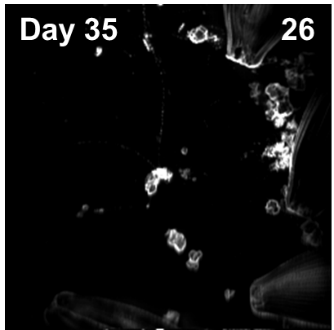

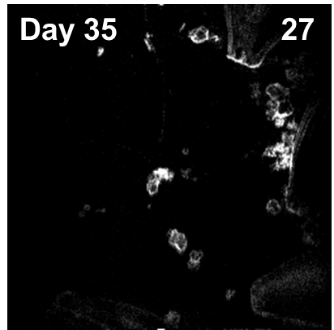

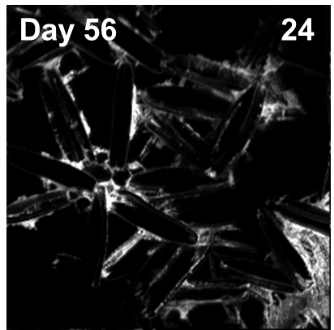

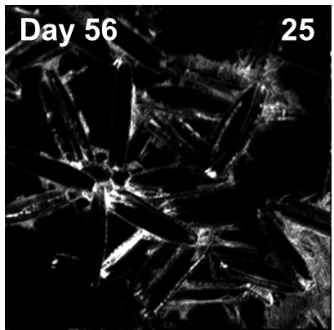

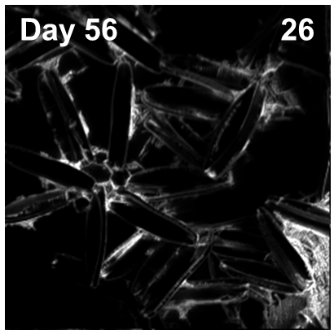

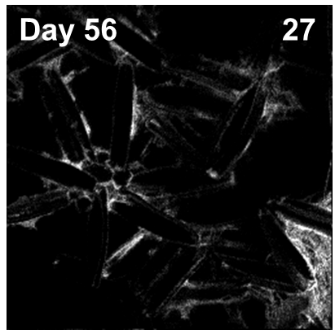

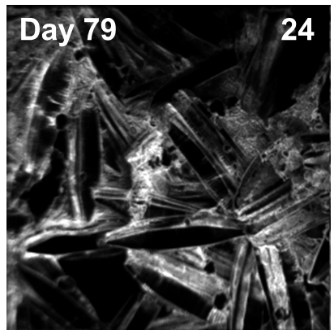

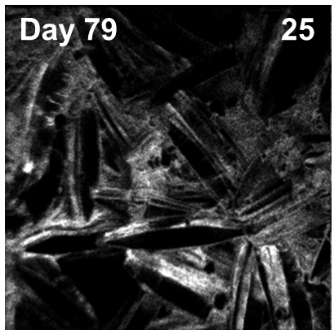

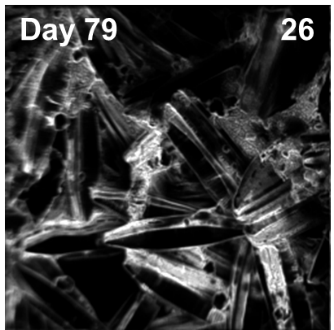

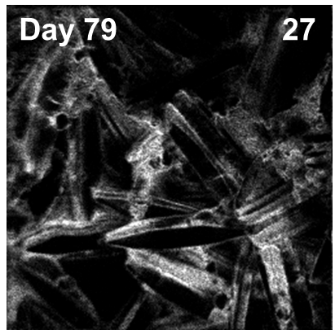

**B) ^15^N-labeled N_2_ incubation**

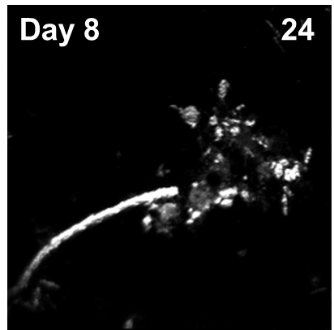

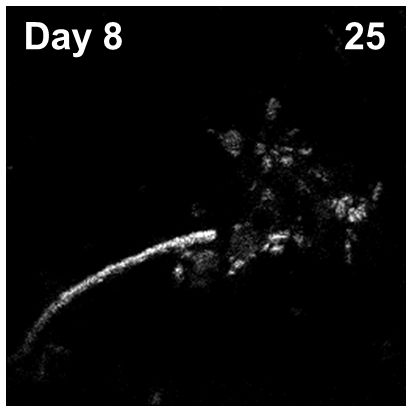

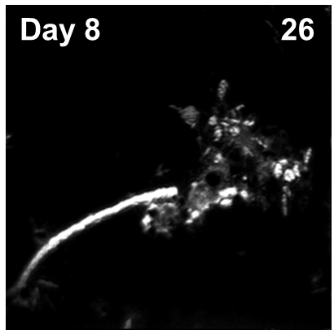

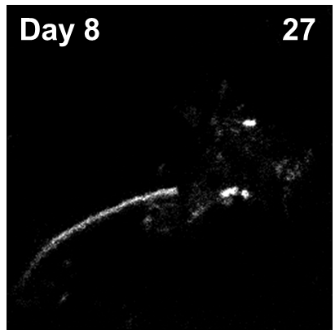

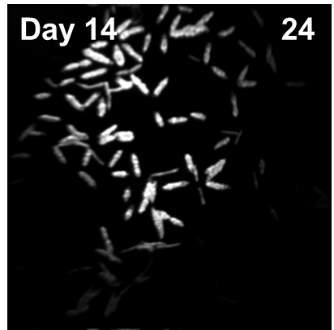

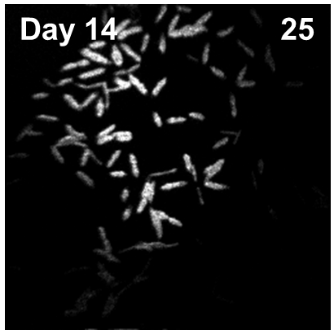

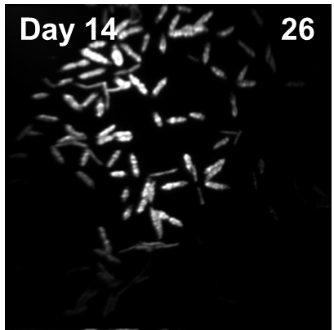

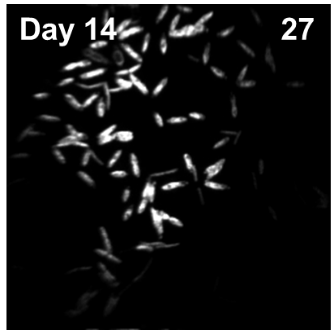

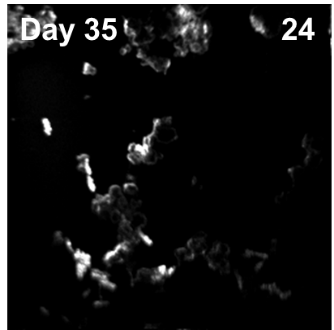

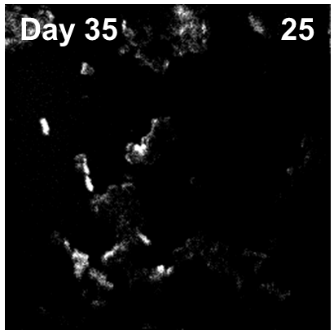

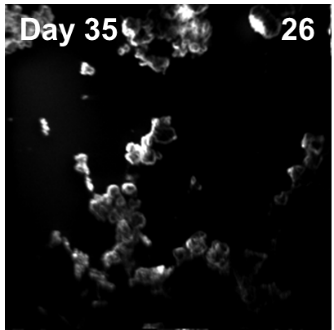

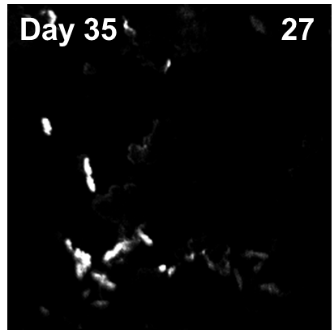

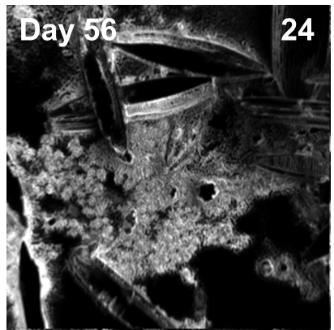

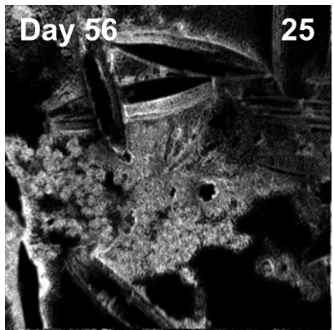

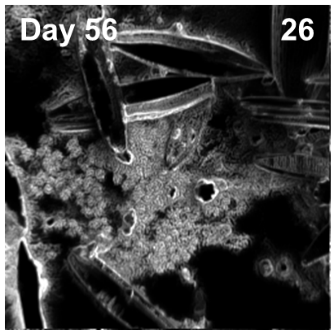

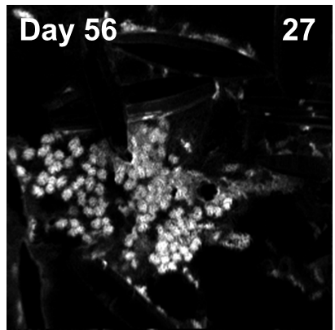

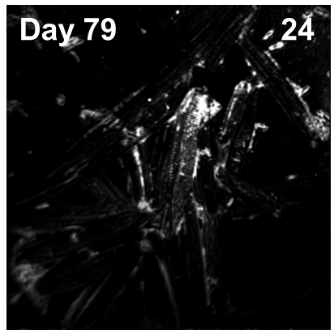

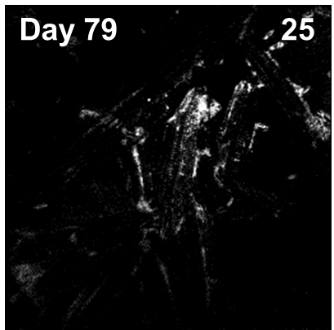

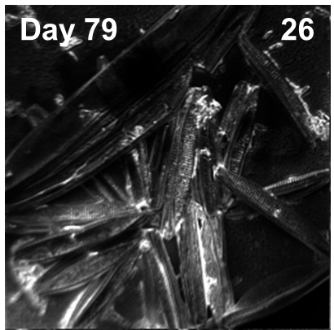

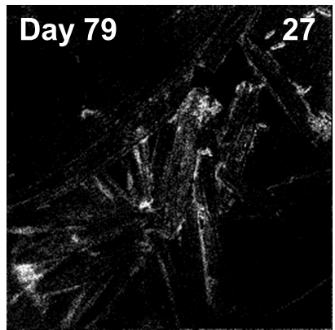

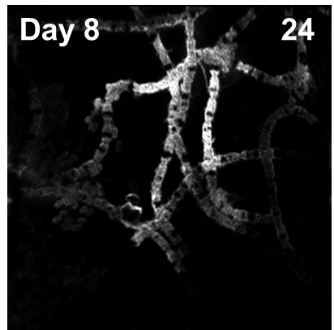

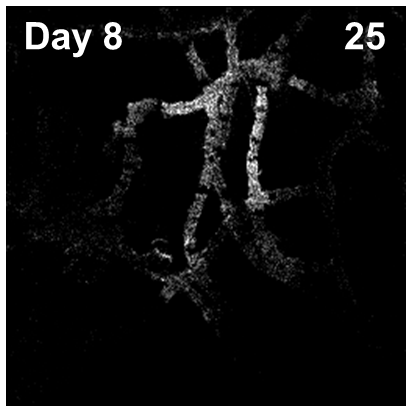

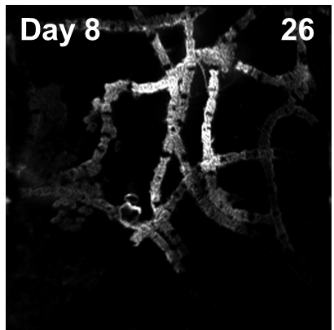

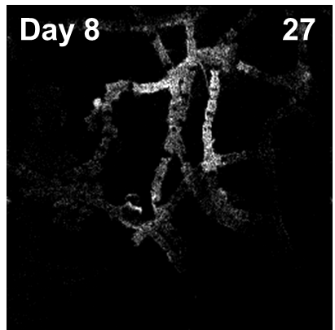

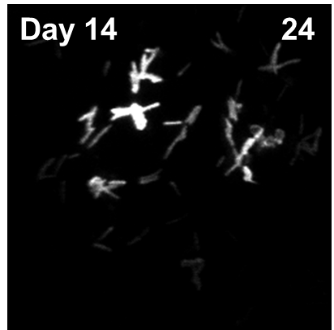

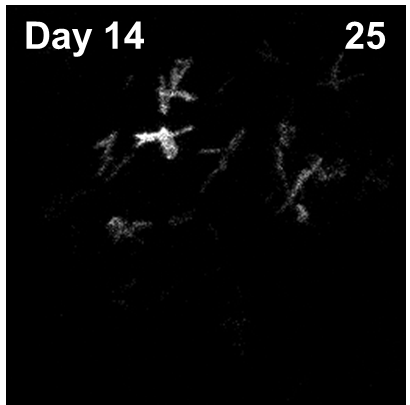

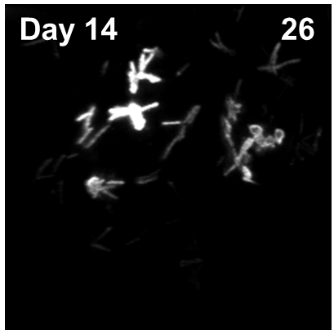

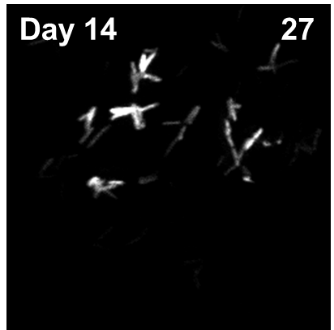

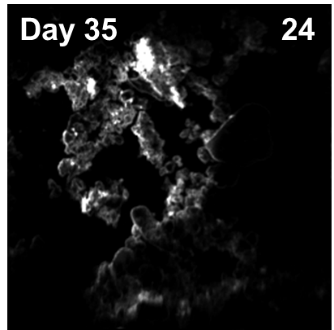

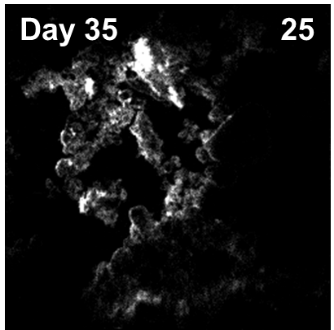

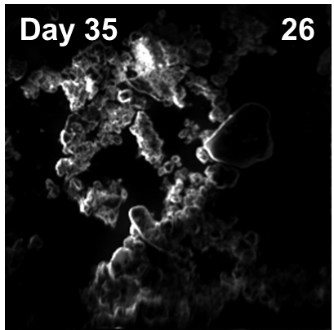

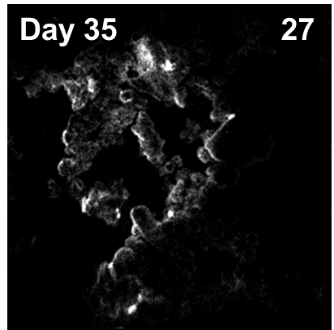

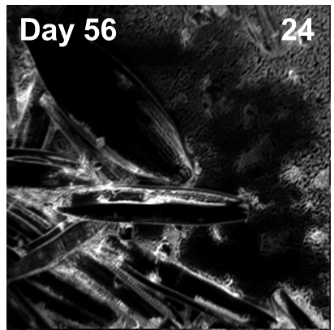

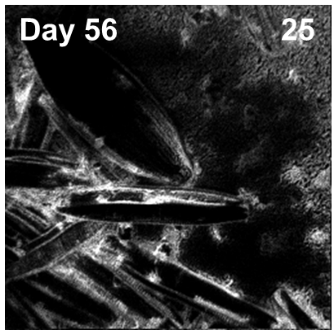

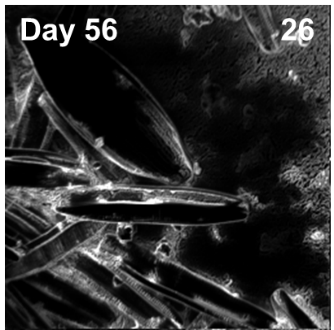

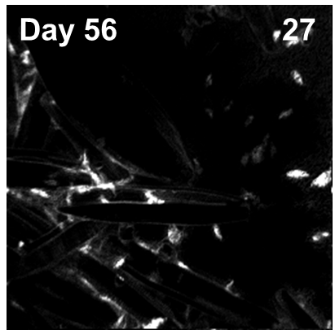

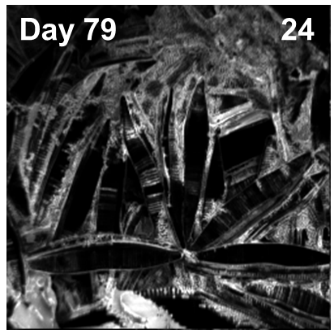

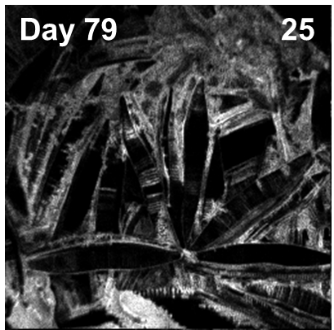

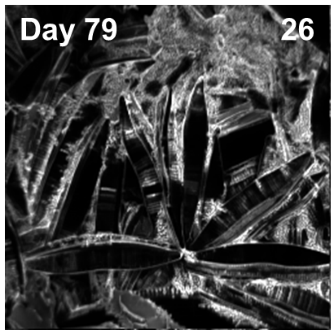

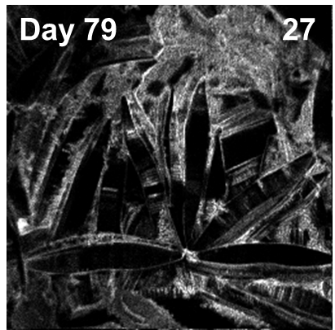

**C) Natural abundance bicarbonate incubation**

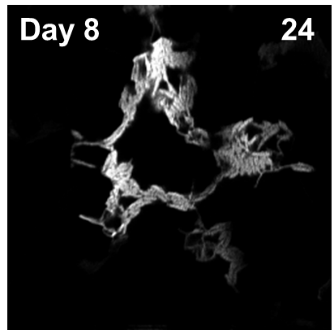

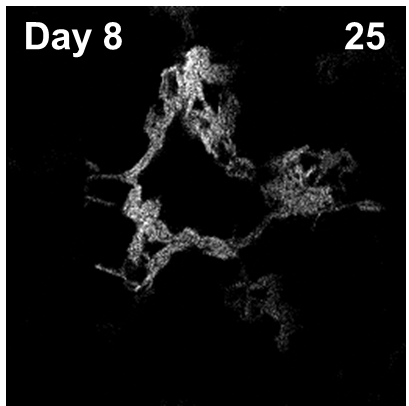

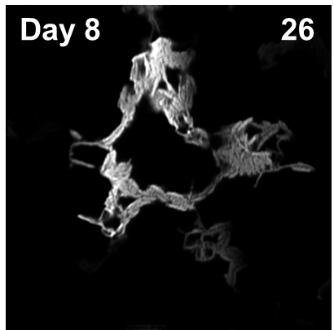

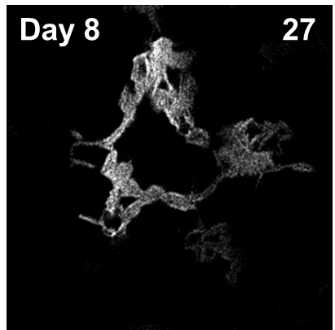

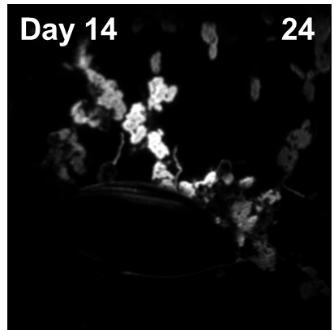

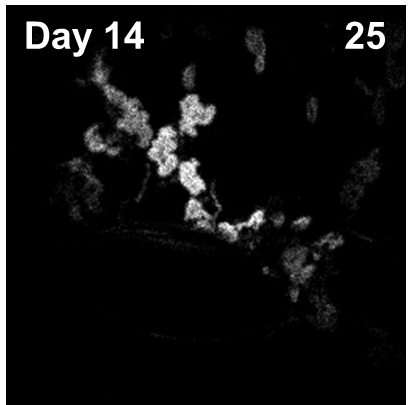

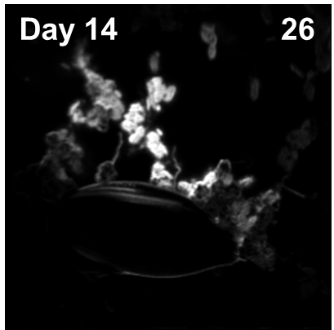

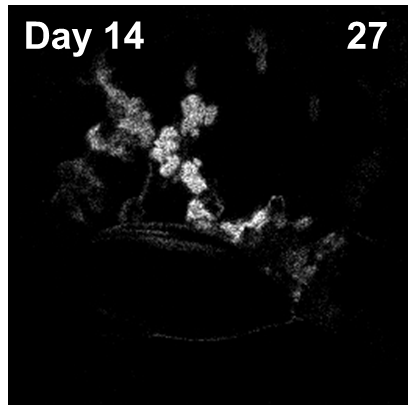

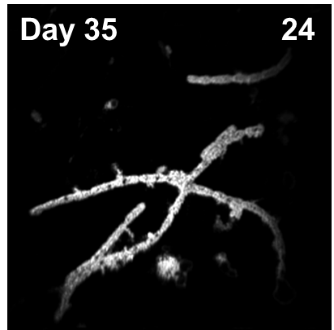

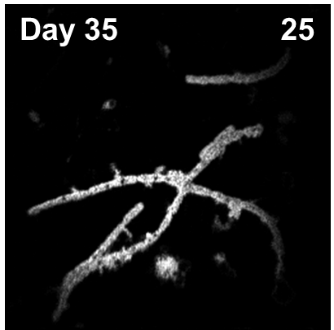

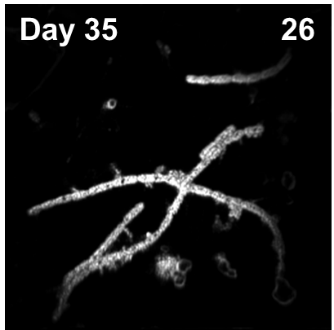

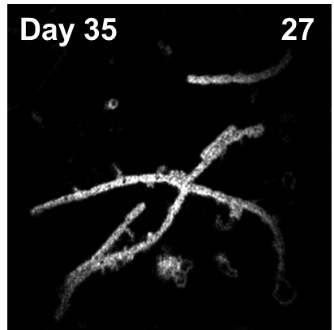

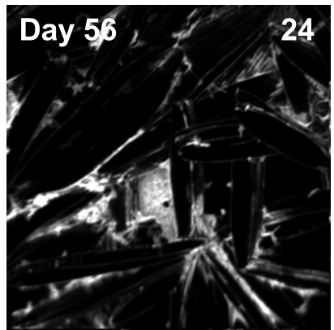

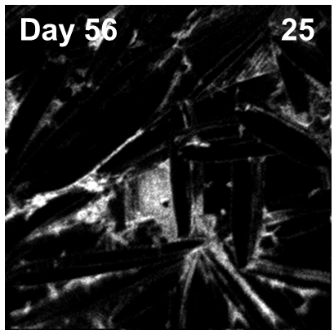

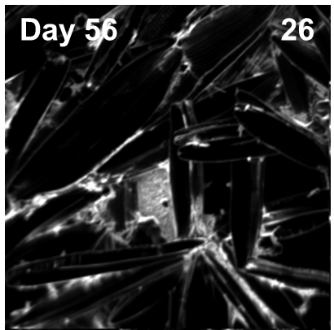

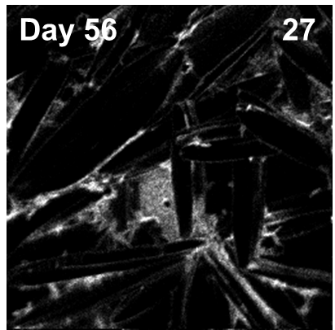

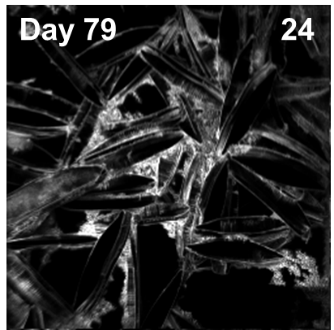

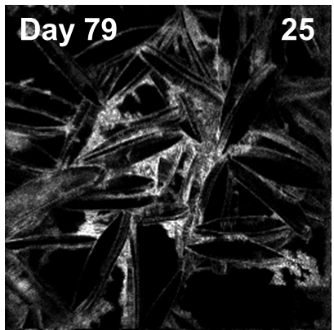

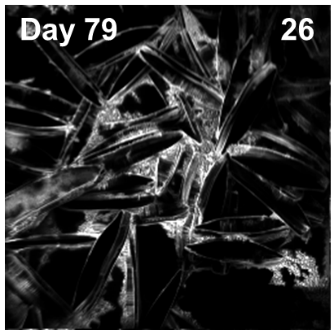

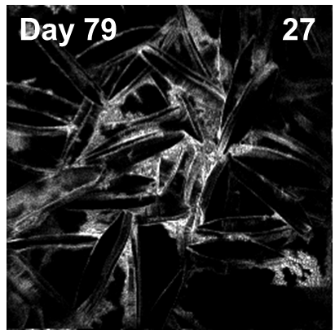

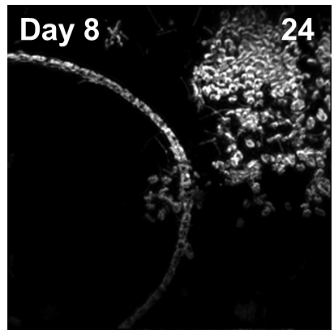

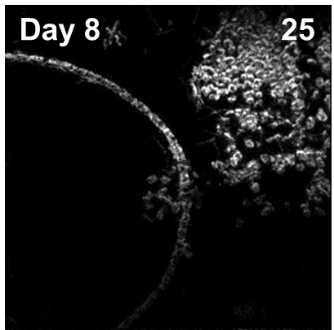

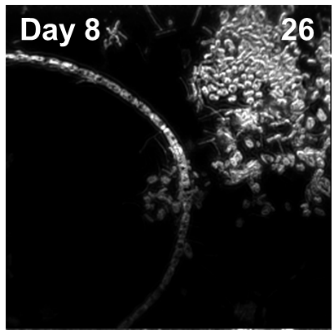

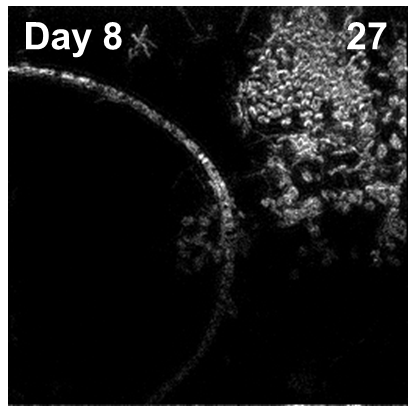

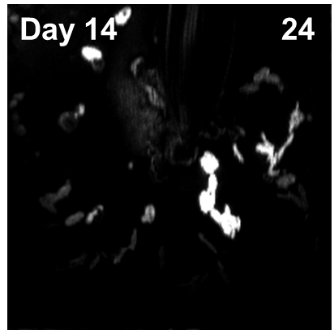

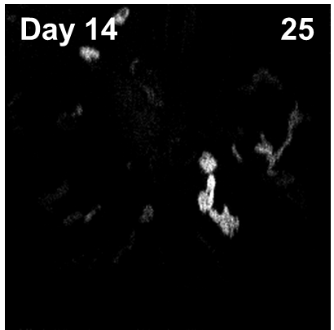

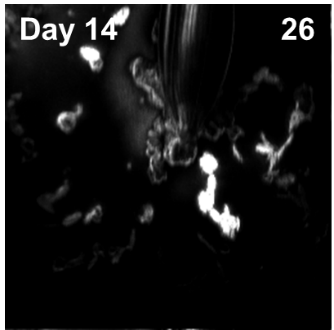

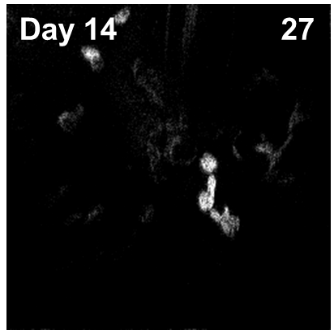

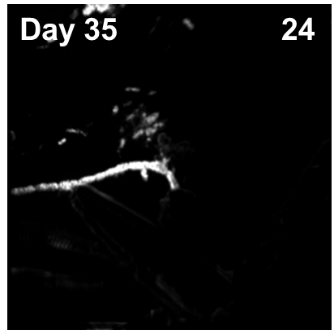

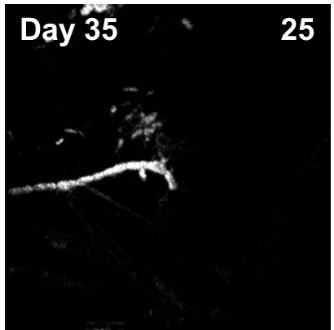

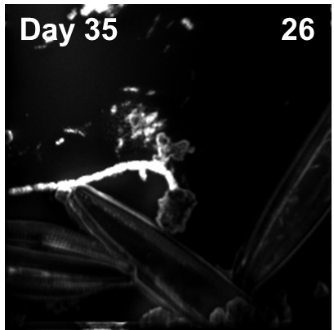

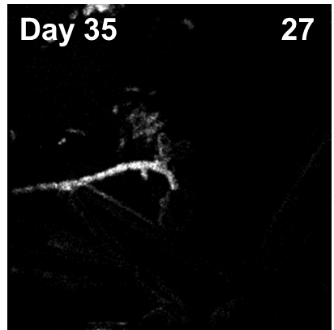

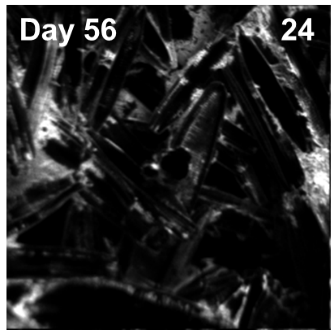

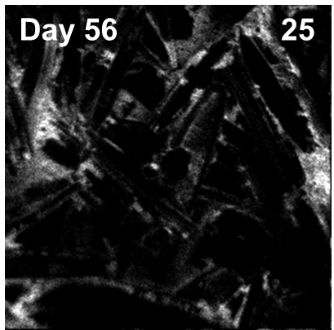

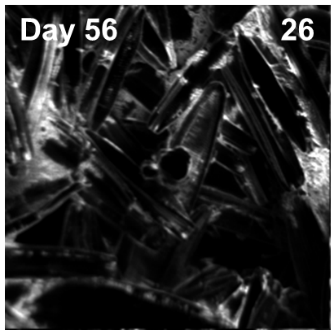

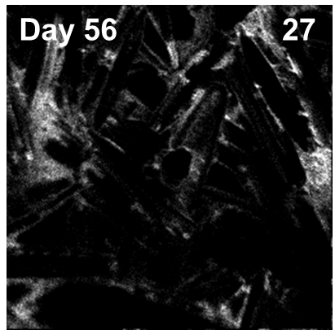

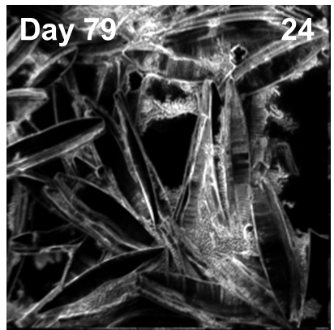

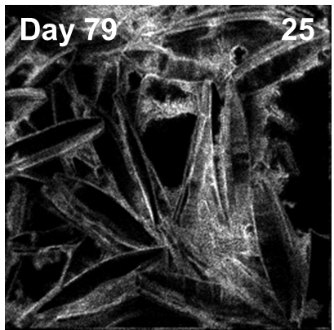

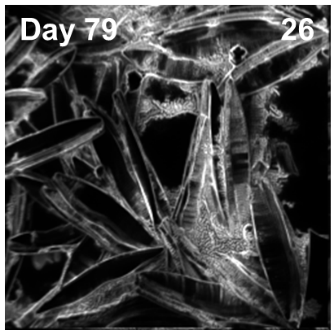

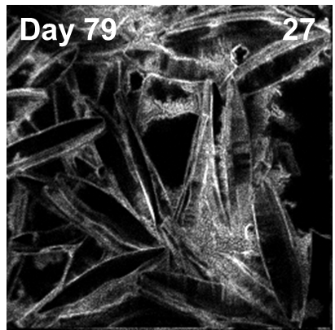

**D) Natural abundance N_2_ incubation**

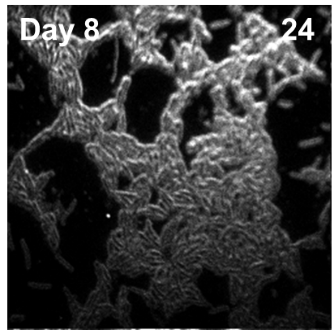

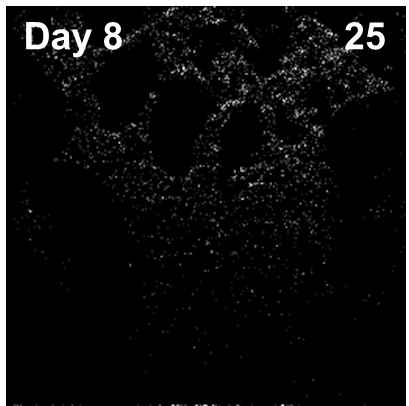

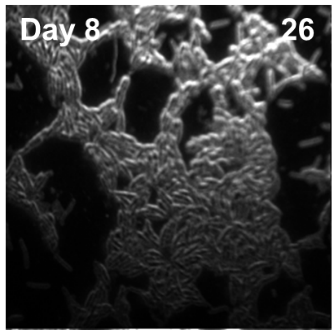

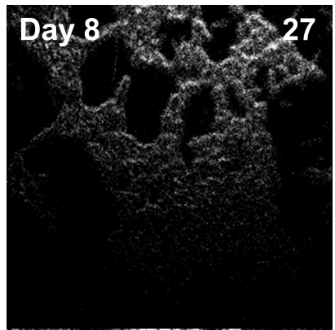

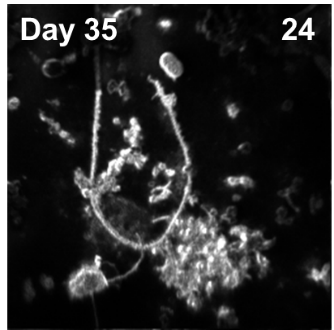

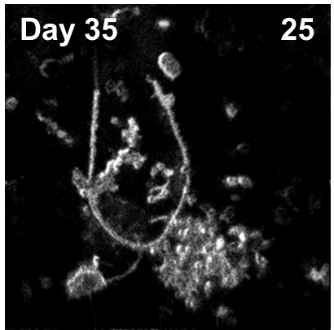

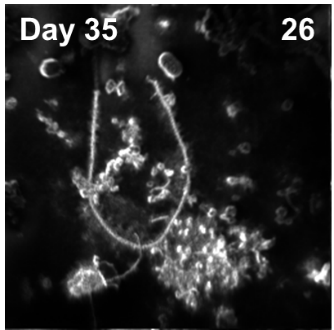

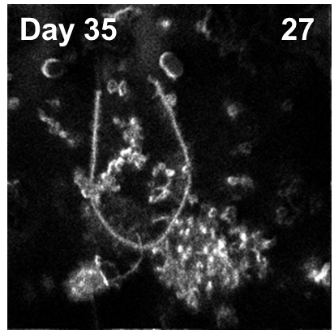

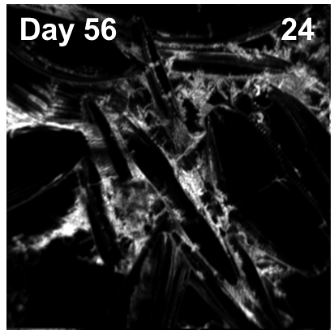

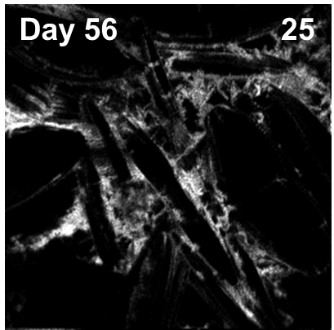

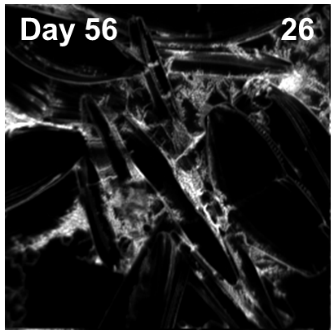

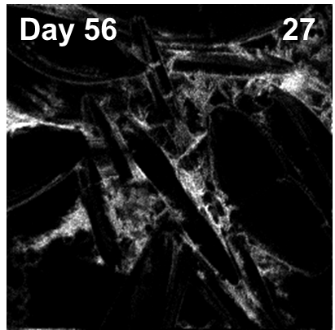

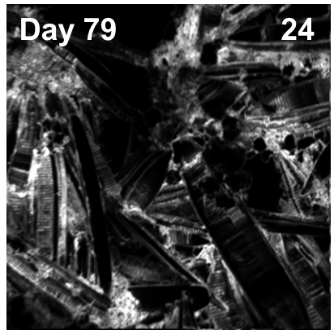

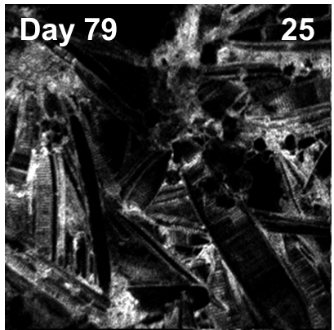

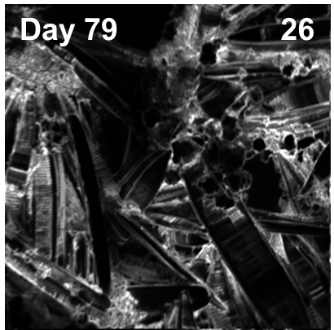

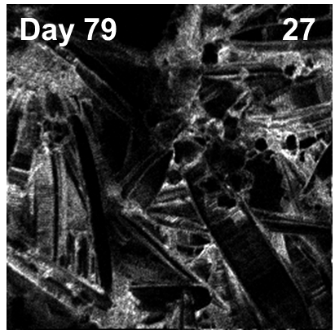

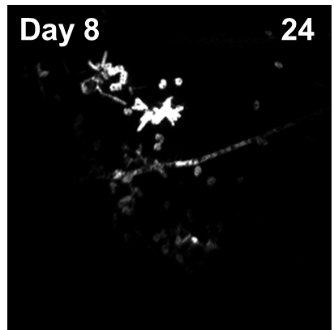

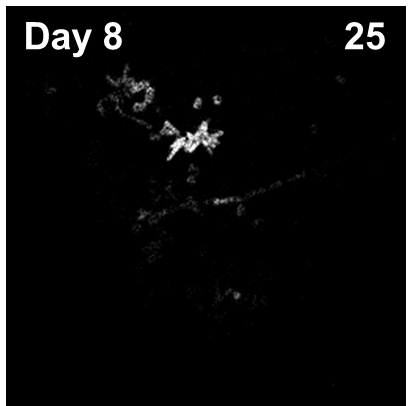

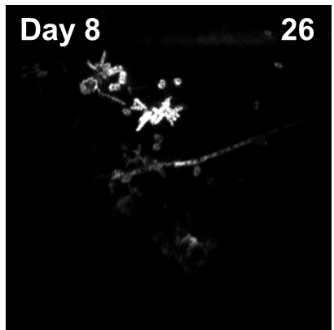

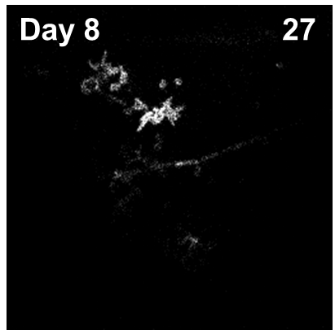

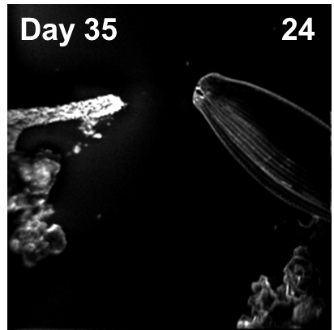

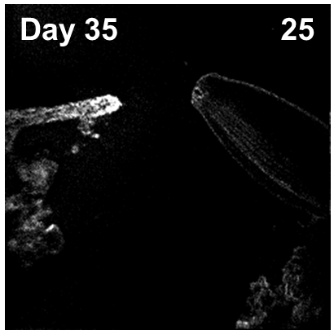

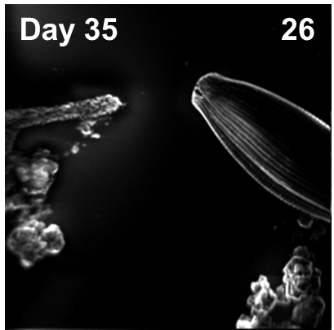

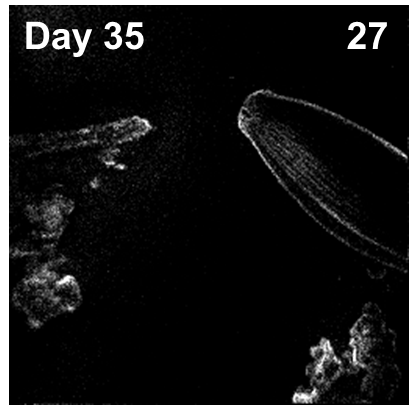

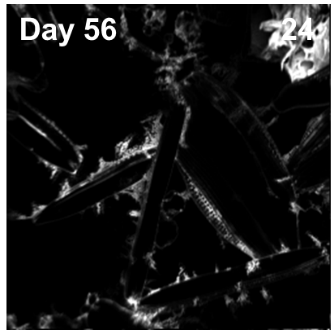

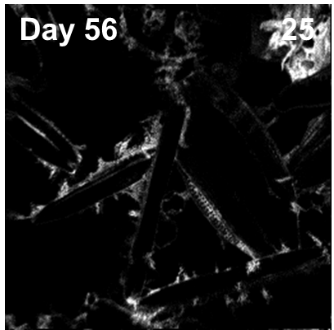

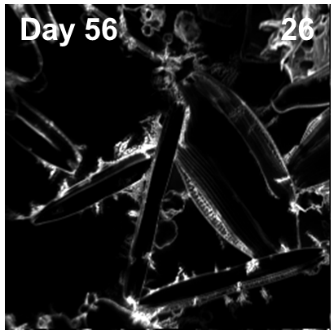

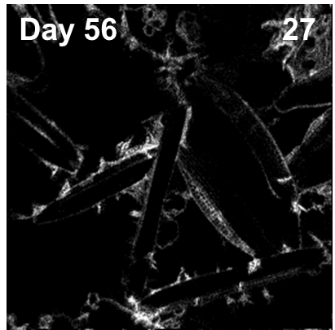

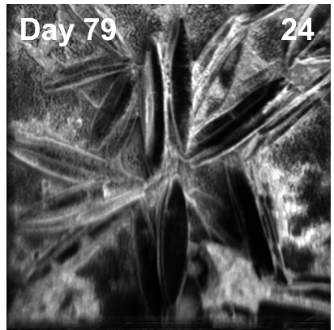

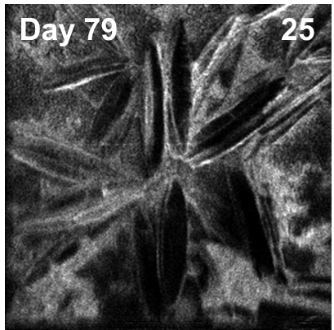

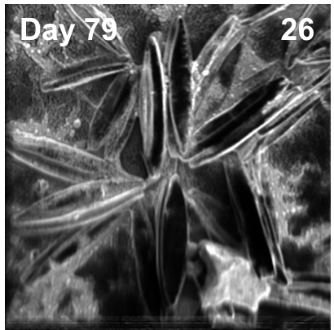

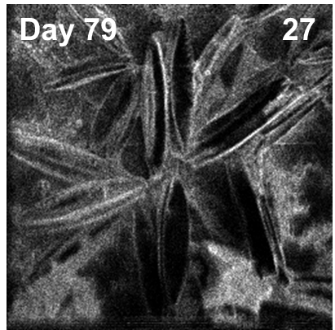

**Figure S3.** NanoSIMS ion intensity images of Hot Lake biofilm samples under different incubation conditions and durations. Image text denotes duration of incubation and m/z (species), where 24 represents ^12^C_2_^-^, 25, ^12^C^13^C^-^, 26, ^12^C^14^N^-^, and 27, ^12^C^15^N^-^. All images are 40 x 40 μm. Replicate analyses shown for each sample type. A: Incubation with ^13^C-labelled bicarbonate. B: Incubation with ^15^N-labelled N_2_. C: Incubation with natural abundance (unlabeled) bicarbonate. D: Incubation with natural abundance (unlabeled) N_2_.

**^13^C-labeled Bicarbonate	^15^N-labeled N_2_	Unlabeled Bicarbonate	Unlabeled N_2_**

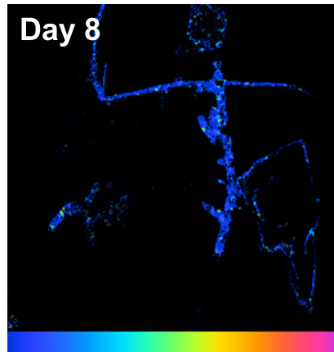

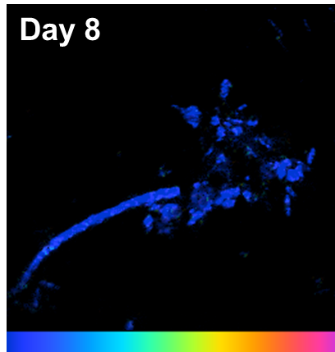

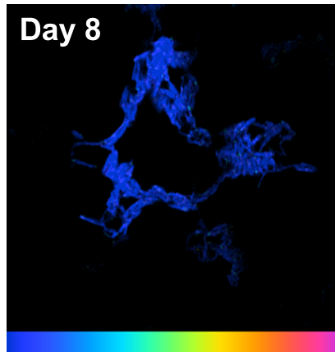

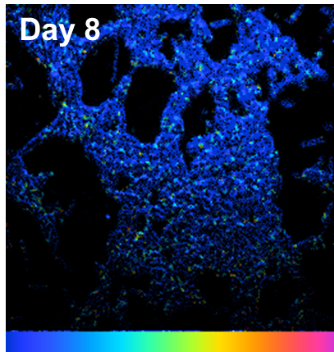

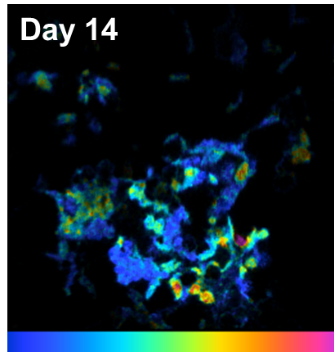

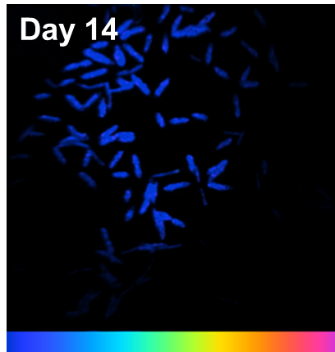

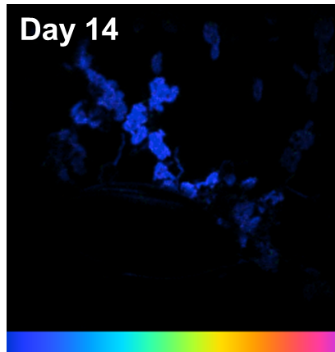

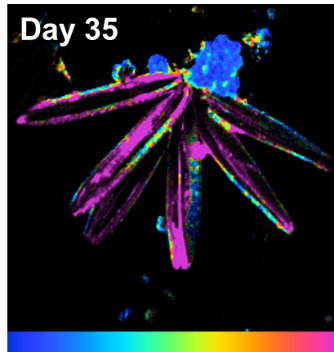

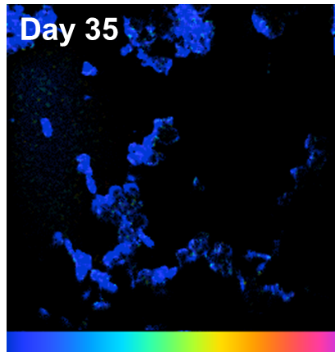

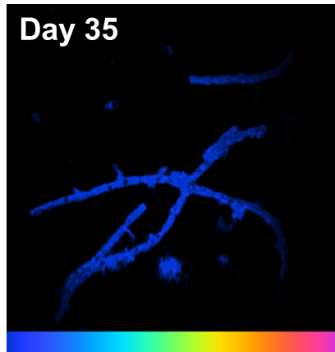

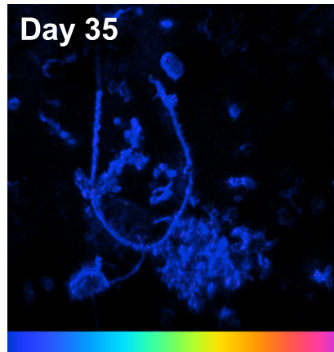

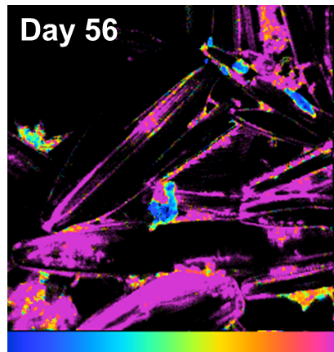

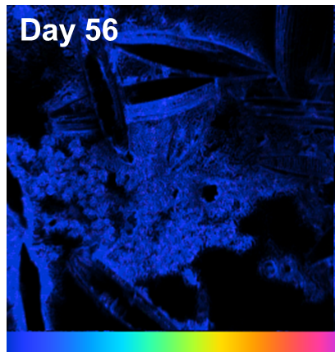

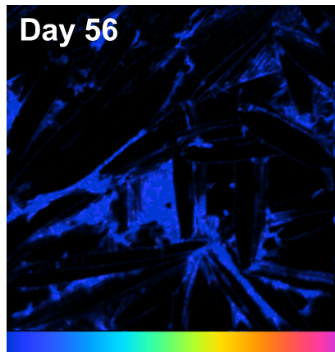

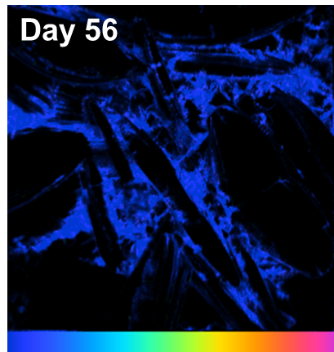

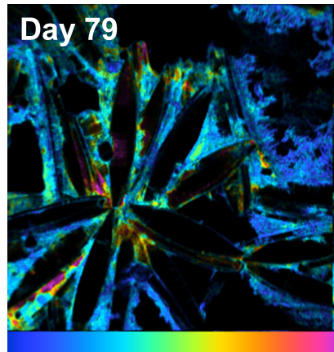

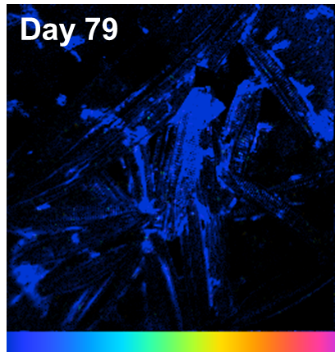

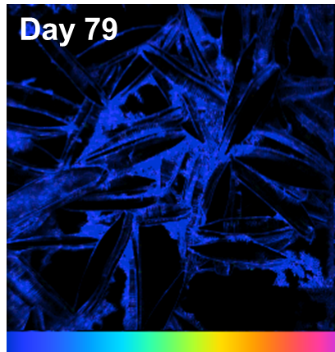

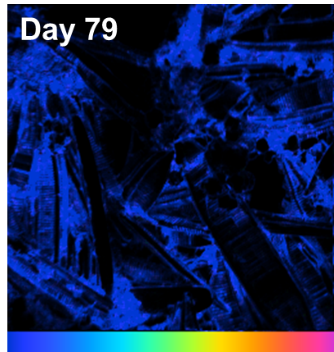

**^13^C-labeled Bicarbonate	^15^N-labeled N_2_	Unlabeled Bicarbonate	Unlabeled N_2_**

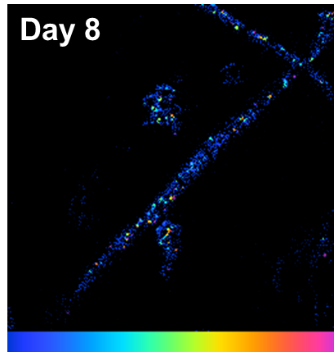

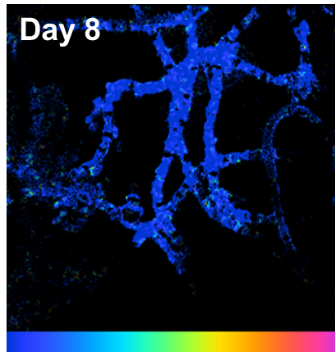

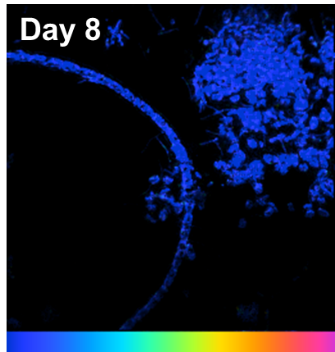

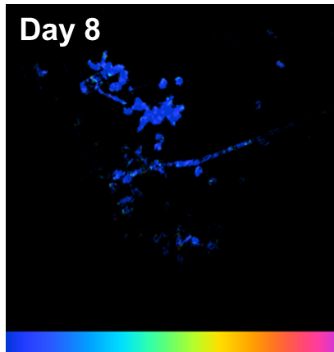

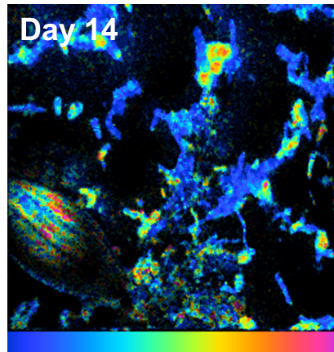

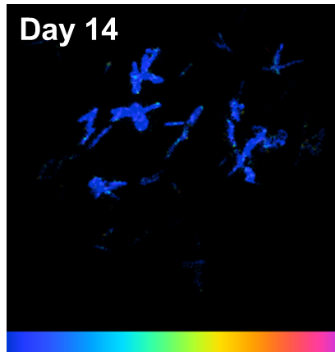

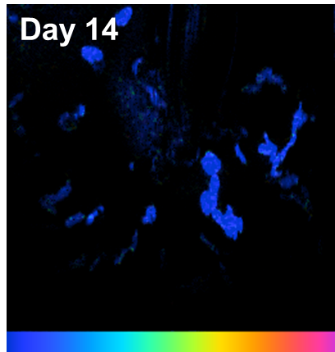

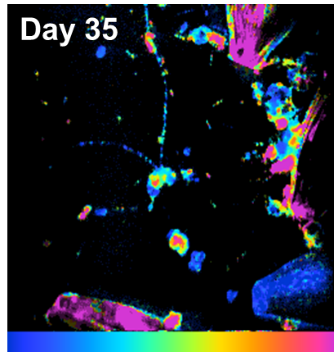

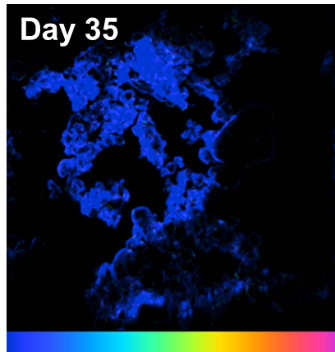

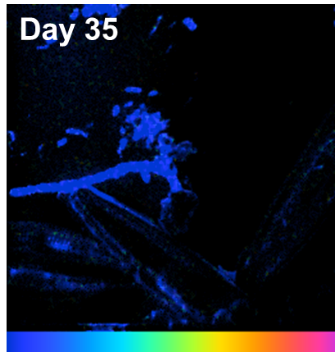

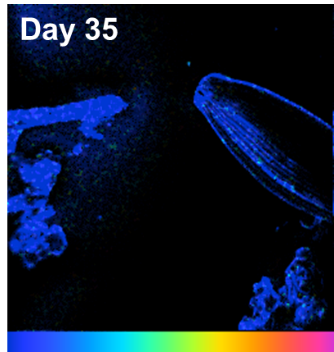

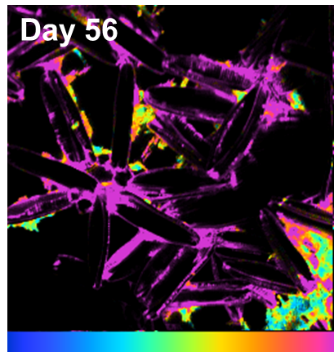

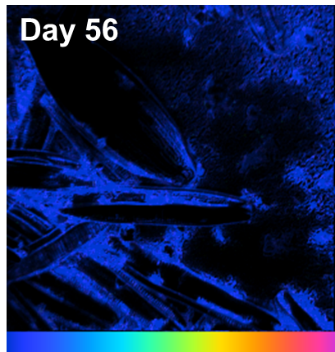

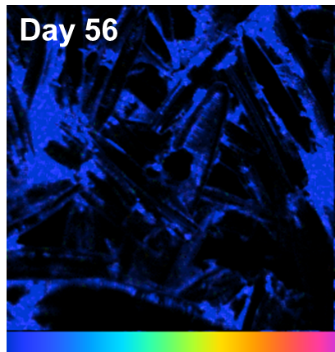

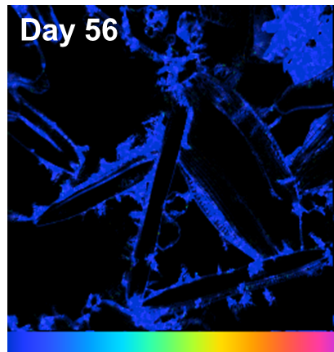

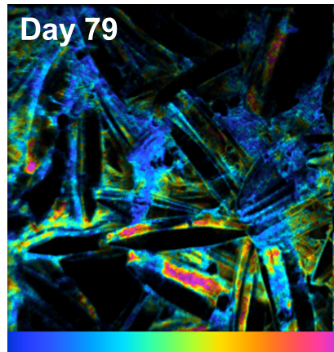

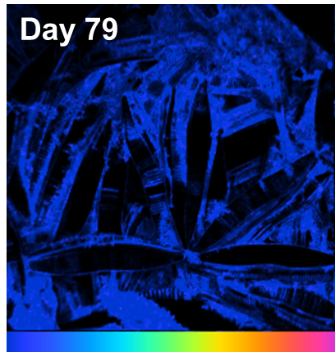

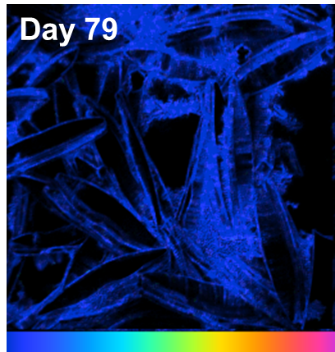

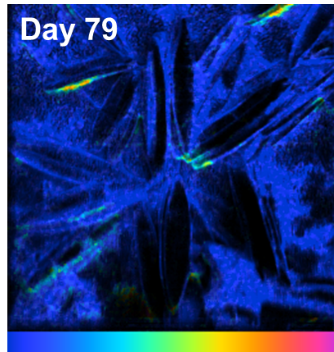

**Figure S4. (above)** NanoSIMS Hue-Saturation-Intensity (HSI) maps of ^13^C^12^C/^12^C_2_ distribution in Hot Lake samples incubated under different conditions and durations. Image text denotes duration of incubation and columns indicate added substrate, as labelled. HSI color range correlates to δ^13^C_PDB_ value, where dark blue (leftmost color on scale bar) is -11‰ and magenta (rightmost color) is >1700‰. Images are 40 x 40 μm. Replicate analyses shown for each sample type.

**^13^C-labeled Bicarbonate	^15^N-labeled N_2_	Unlabeled Bicarbonate	Unlabeled N_2_**

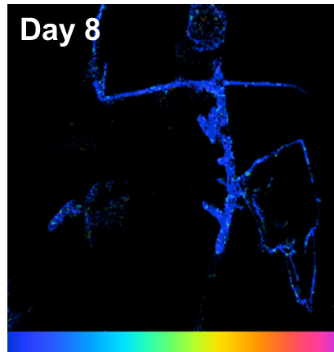

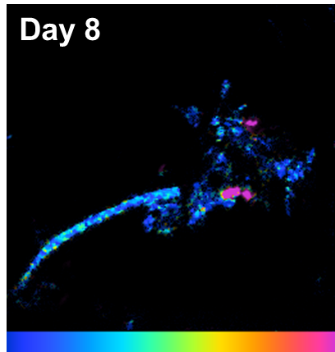

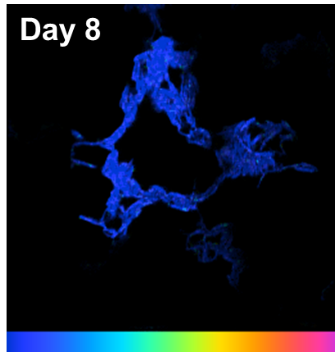

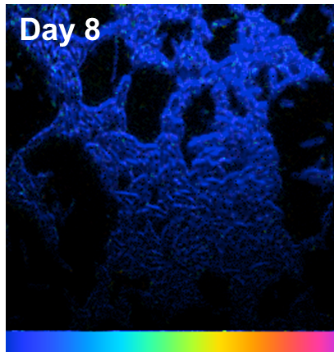

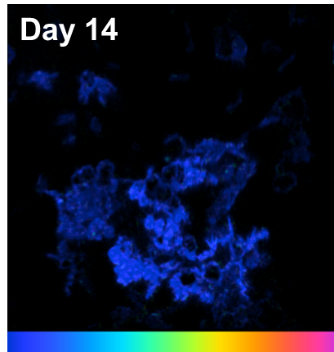

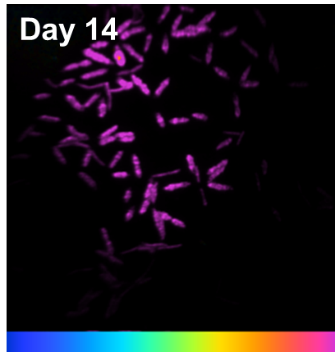

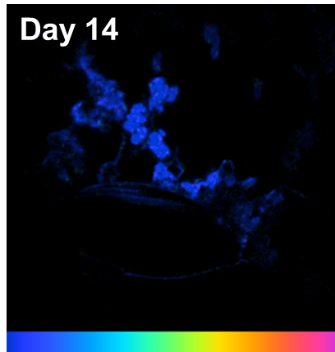

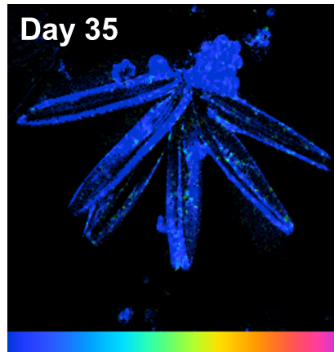

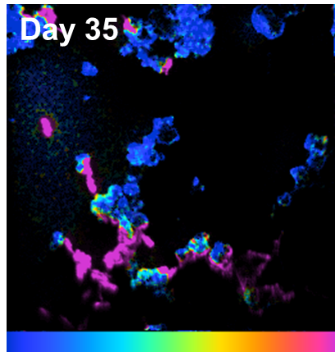

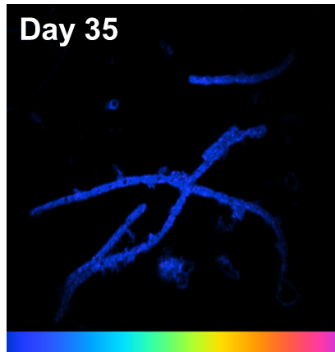

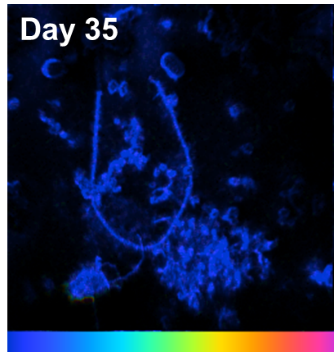

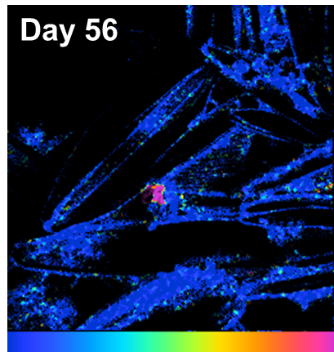

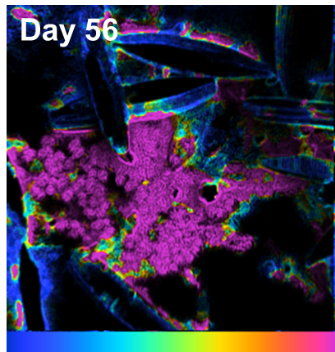

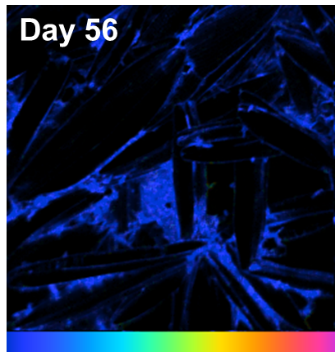

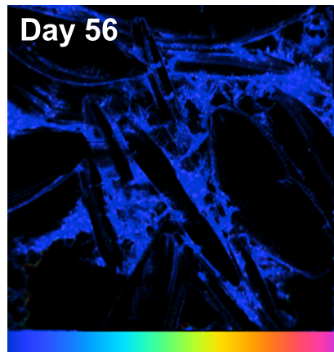

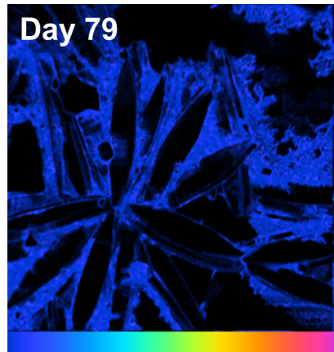

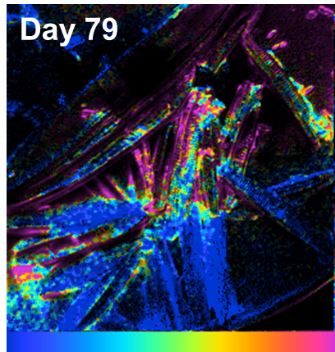

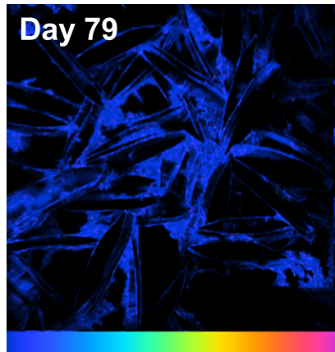

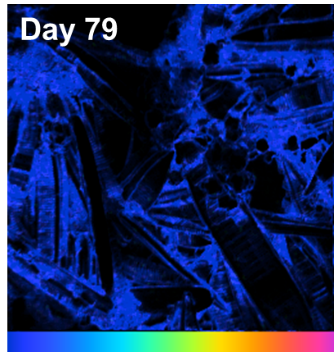

**^13^C-labeled Bicarbonate	^15^N-labeled N_2_	Unlabeled Bicarbonate	Unlabeled N_2_**

**^13^C-labeled Bicarbonate	^15^N-labeled N_2_	Unlabeled Bicarbonate	Unlabeled N_2_**

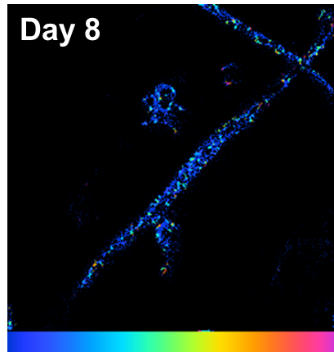

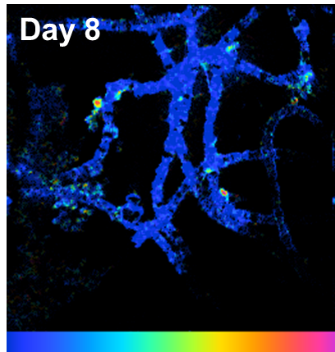

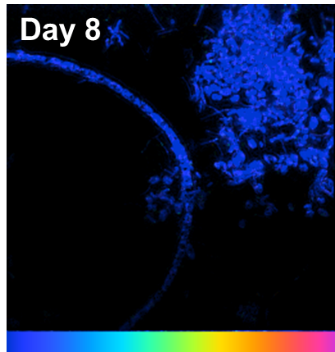

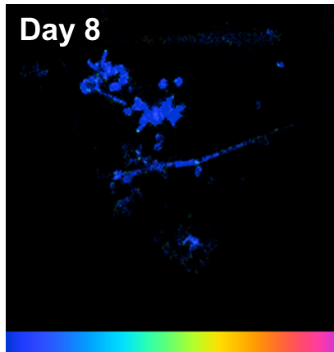

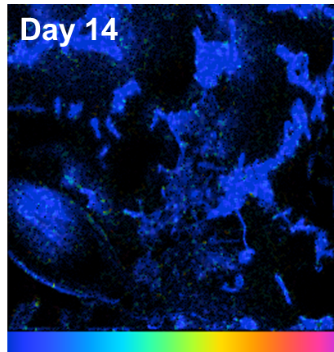

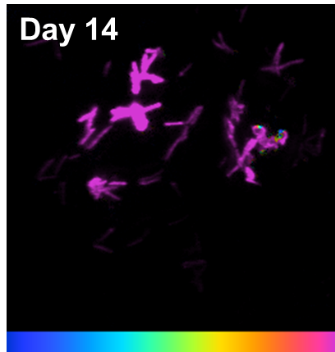

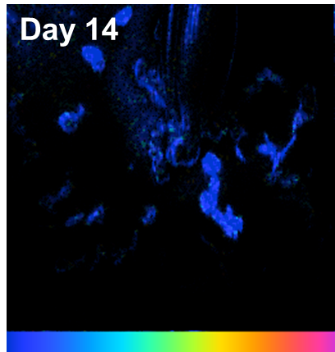

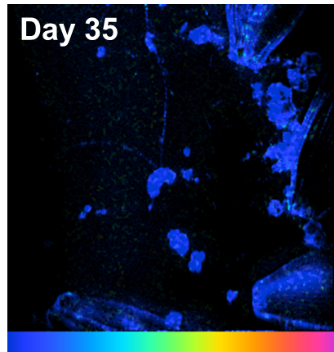

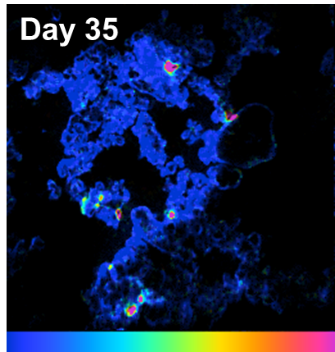

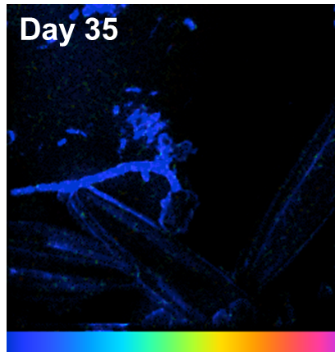

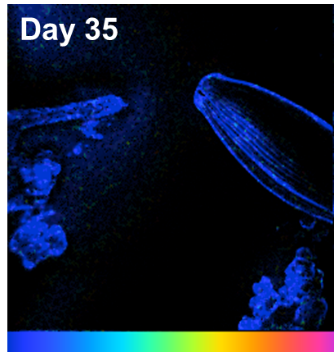

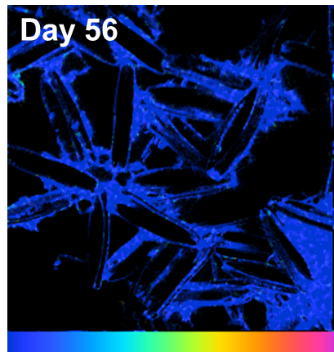

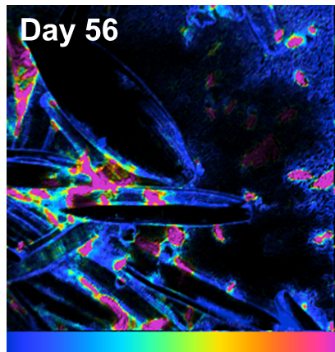

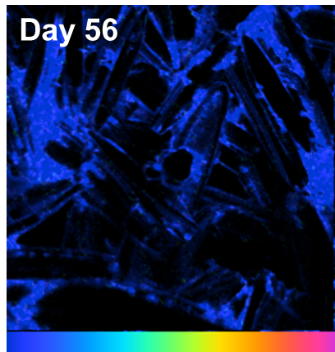

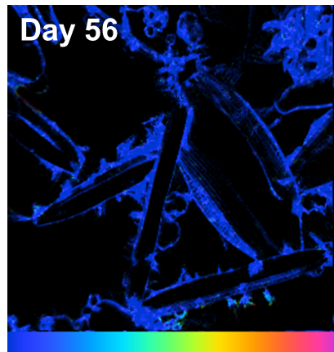

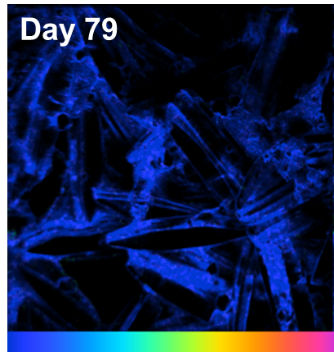

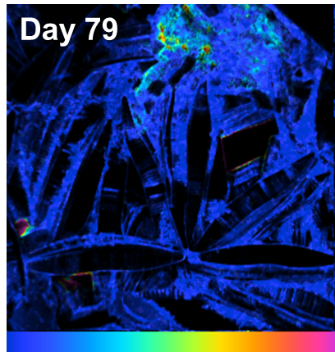

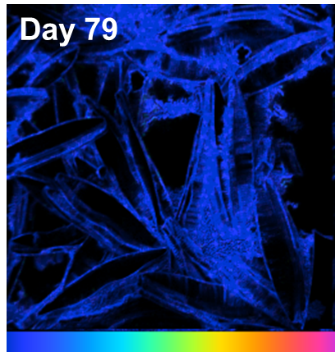

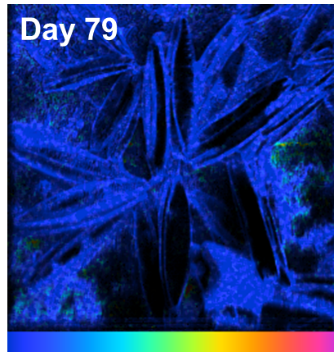

**^13^C-labeled Bicarbonate	^15^N-labeled N_2_	Unlabeled Bicarbonate	Unlabeled N_2_**

**Figure S5.** NanoSIMS Hue-Saturation-Intensity (HSI) maps of ^12^C^15^N/^12^C^14^N distribution in Hot Lake samples incubated under different conditions and durations. Image text denotes duration of incubation and columns indicate added substrate, as labelled. HSI color range correlates to δ^15^N_AIR_ value, where dark blue (leftmost color on scale bar) is 0‰ and magenta (rightmost color) is >1700‰. Images are 40 x 40 μm. Replicate analyses shown for each sample type.

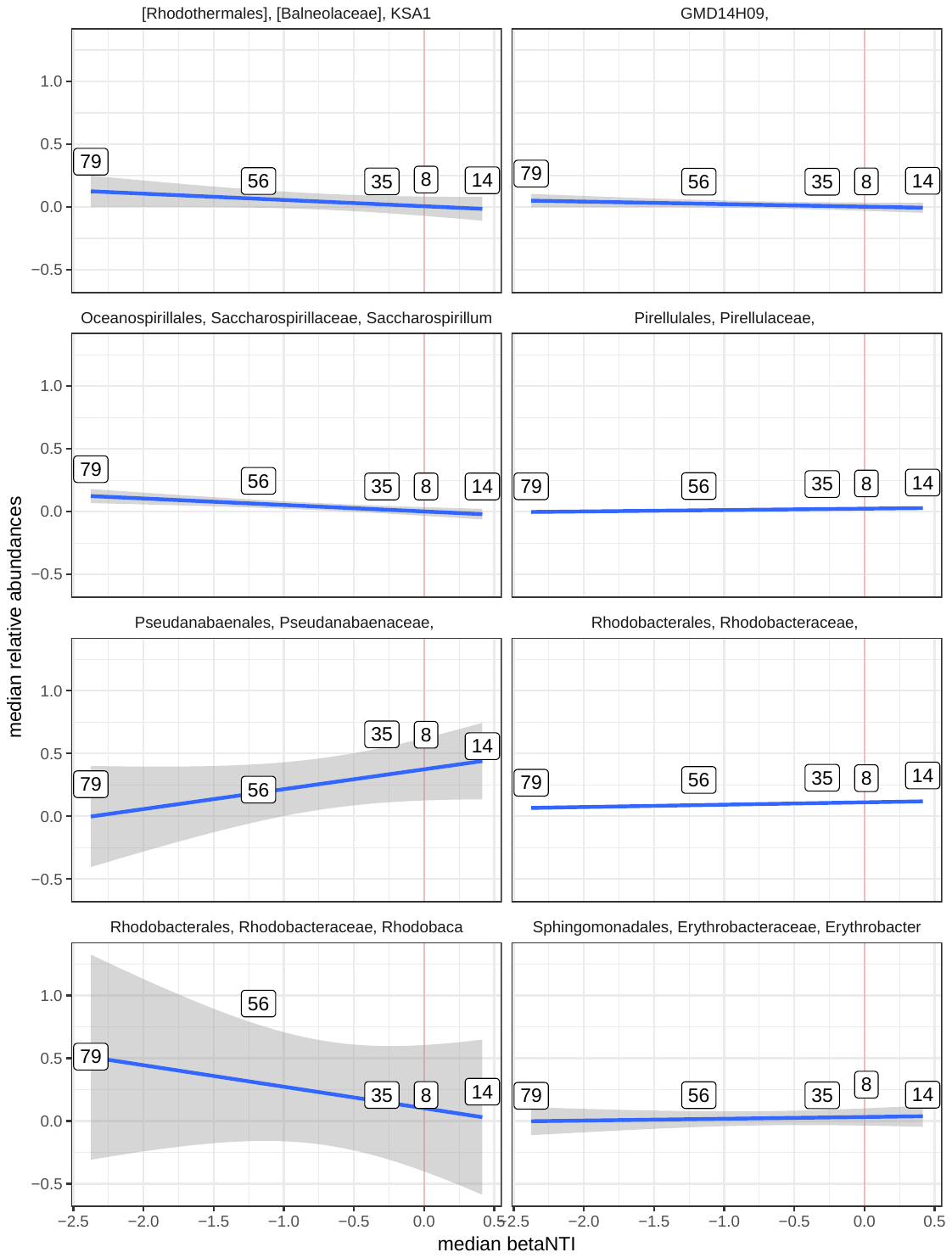

**Figure S6. Correlation between bacterial founder species and selective pressure over time.** Between all pairs of samples at each day, the median βNTI was calculated in order to measure if samples were under significant homogeneous (βNTI < -2) or heterogeneous (βNTI > +2) selective pressure.

This is compared to the median relative abundance of each founder species at that day. A positive correlation between abundance and Beta NTI shows that a taxon has higher relative abundances during variable selection as observed on early days (i.e., Pseudanabaenaceae). Anti-correlation indicates that a taxon thrives under homogeneous, stabilizing selection, as observed on later days (i.e., Rhodobaca).

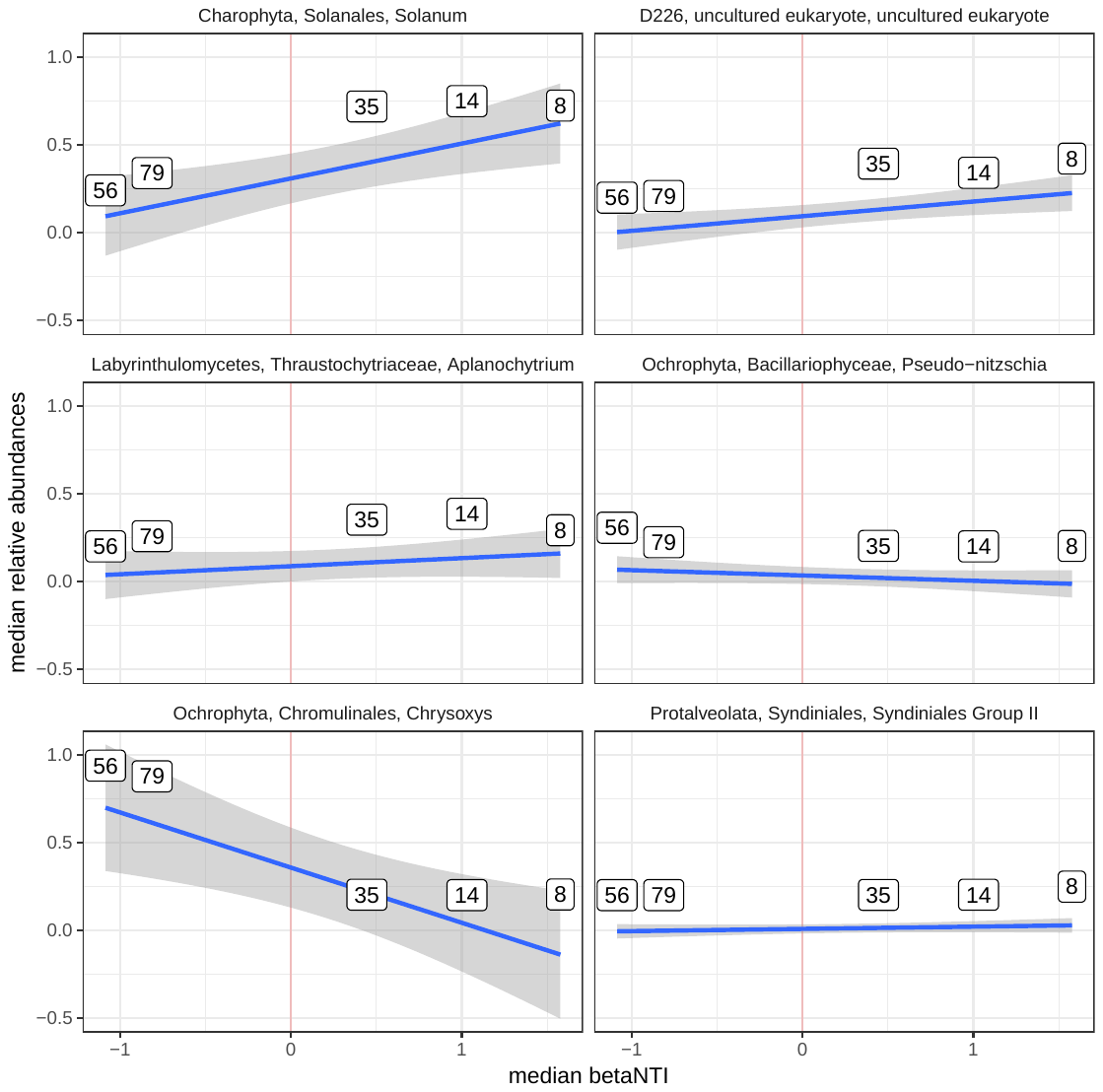

**Figure S7.** Beta NTI within each time point for eukaryotic founder species

**Table S1.** The adonis test is used to partition variation of nestedness and turnover by sampling day. The partial R-squared value shows what fraction of variation of binary Jaccard distances can be explained by Day, and p-values are calculated based on 999 permutations.

| **Microbe** | **Type** | **Df** | **SumsOfSqs** | **MeanSqs** | **F.Model** | **R2** | **p-value** |
| --- | --- | --- | --- | --- | --- | --- | --- |
| Bacteria | Total | 4 | 5.7138837 | 1.4284709 | 8.3450911 | 0.3358849 | 0.001 |
| Bacteria | Turnover | 4 | 1.8046338 | 0.4511584 | 4.1079679 | 0.1993388 | 0.001 |
| Bacteria | Nestedness | 4 | 0.8394003 | 0.2098501 | 17.0852862 | 0.5087134 | 0.001 |
| Eukaryotes | Total | 4 | 4.3456018 | 1.0864005 | 5.0363380 | 0.2365995 | 0.001 |
| Eukaryotes | Turnover | 4 | 3.2097125 | 0.8024281 | 5.8863480 | 0.2659132 | 0.001 |
| Eukaryotes | Nestedness | 4 | -0.0061397 | -0.0015349 | -0.1099199 | -0.0068104 | 0.985 |
| Combined | Total | 4 | 4.8632051 | 1.2158013 | 4.6864211 | 0.1898380 | 0.001 |
| Combined | Turnover | 4 | 3.3103050 | 0.8275763 | 4.7370747 | 0.1914970 | 0.001 |
| Combined | Nestedness | 4 | 0.2188416 | 0.0547104 | 3.5033644 | 0.1490580 | 0.004 |

**Table S2. Correlation between bacterial founder species and selective pressure over time.** As shown in Fig. S6, the median daily Beta NTI was correlated with the median relative abundance of each founder species at that day. A positive or negative slope of the linear model indicates that taxa thrive under heterogeneous or homogeneous selective pressure, respectively, and the p value reports if this relationship is significantly different from zero.

| **Taxonomy** | **estimate** | **std.error** | **statistic** | **p.value** |
| --- | --- | --- | --- | --- |
| [Rhodothermales], [Balneolaceae], KSA1 | -0.0520451 | 0.0189346 | -2.7486761 | 0.0708208 |
| GMD14H09, | -0.0211188 | 0.0081286 | -2.5980996 | 0.0805079 |
| Oceanospirillales, Saccharospirillaceae, Saccharospirillum | -0.0531663 | 0.0068511 | -7.7602492 | 0.0044512 |
| Pirellulales, Pirellulaceae, | 0.0109994 | 0.0026370 | 4.1711691 | 0.0250851 |
| Pseudanabaenales, Pseudanabaenaceae, | 0.1646620 | 0.0611066 | 2.6946693 | 0.0741200 |
| Rhodobacterales, Rhodobacteraceae, | 0.0188475 | 0.0029819 | 6.3206153 | 0.0080053 |
| Rhodobacterales, Rhodobacteraceae, Rhodobaca | -0.1749548 | 0.1314402 | -1.3310602 | 0.2752846 |
| Sphingomonadales, Erythrobacteraceae, Erythrobacter | 0.0139280 | 0.0181183 | 0.7687276 | 0.4980290 |

**Table S3.** Analogous statistics from Table S2 but with respect to the eukaryotic OTU assignments and regressions shown in Figure S7.

| **Taxonomy** | **estimate** | **std.error** | **statistic** | **p.value** |
| --- | --- | --- | --- | --- |
| Charophyta, Pinales, Pinus | 0.0067142 | 0.0043864 | 1.530691 | 0.2233448 |
| Chlorophyta, Chlamydomonadales, Dunaliella | 0.0081180 | 0.0028658 | 2.832751 | 0.0660397 |
| D226, uncultured eukaryote, uncultured eukaryote | 0.1512910 | 0.0287425 | 5.263666 | 0.0133619 |
| Labyrinthulomycetes, Thraustochytriaceae, Aplanochytrium | 0.0902216 | 0.0563124 | 1.602164 | 0.2074423 |
| Ochrophyta, Bacillariophyceae, Pseudo-nitzschia | -0.0291968 | 0.0177864 | -1.641525 | 0.1992269 |
| Ochrophyta, Chromulinales, Chrysoxys | -0.2999766 | 0.0559553 | -5.361005 | 0.0127011 |
| Protalveolata, Syndiniales, Syndiniales Group II | 0.0287754 | 0.0172041 | 1.672590 | 0.1930014 |
